## Supplemental Information for "Specificity, synergy, and mechanisms of splice-modifying drugs"

### *Specificity, synergy, and mechanisms of splice-modifying drugs: Supplemental Information*

#### *1 Experimental Methods*

##### 1.1 Cell culture

HeLa cells were obtained from the Cold Spring Harbor Laboratory Cell Line Repository shared resource, which validates cell lines by genotyping and routinely checks for Mycoplasma contamination. HeLa cells were seeded at $2\times10^6$ on 15-cm plates and cultured for 72 hrs at 37 ˚C, 5% $CO_2$. Cells were then re-seeded at $2.5\times10^5$ cells on 12-well plates and cultured for 16-24 hrs. Cells were then transfected by a mixture of (i) 5 µl of Lipofectamine 2000 in 62.5 µl of Opti-MEM and (ii) 2 µg minigene plasmid, plus 2.5 µl of water with or without 1000x ASO in 62.5 µl Opti-MEM. Immediately after transfection, 5 µl of DMSO (plus or minus 500x risdiplam, 500x branaplam, or 500x RECTAS) in 115 µl DMEM was added to the wells and mixed. After transfection and the addition of drugs, cells were cultured for an additional 48 hrs.

##### 1.2 RNA extraction

Cells were washed with PBS. Total RNA was extracted by adding 750 µl TRIzol, then adding 150 µl chloroform and vortexing, then performing centrifugation at 12,000g for 10-20 min. 300 µl of supernatant was then added to 300 µl of iPrOH and centrifuged at 20,000g for 30 min. The supernatant was removed, the pellet was rinsed with 75% EtOH, and RNA was resuspended in nuclease-free water. TURBO DNase treatment was applied to 5.5 µg of total RNA according to the manufacturer's protocol.

##### 1.3 Minigene plasmids

As in ref.[1], all minigenes were cloned into a pcDNA5 plasmid backbone, were expressed from a CMV promoter, and were terminated by a BGH polyadenylation signal. All minigene plasmids used in this study are listed in **Table S2**. SnapGene files for plasmids pSMN2_WT and pELP1_WT are provided on GitHub. These SnapGene files include annotations describing all the plasmid variants listed in **Table S2**.

Minigene plasmid pSMN2_WT contains an *SMN2* minigene spanning exons 6, 7, and 8. Specifically, the minigene contains all of exon 6 (111 nt), part of intron 6 (1393 nt, reduced from 5769), all of exon 7 (54 nt), all of intron 7 (444 nt), and part of exon 8 (first 75 nt). This minigene was formed from two genomic segments joined within intron 6 at position +61.5. Segment 1 (173 nt) derives from ch5:70,707,640..70,070,812 (GRCh38 coordinates), and segment 2 (1905 nt) derives from ch5:70,075,191..70,077,093. Intron 6 has three additional mutations in segment 2: Δ+103..+104, C+141T, and C-376A. Intron 7 has an additional mutation, T+25A, which removes a cryptic 5'ss.

Multiple variants of pSMN2_WT were constructed. In pSMN2_cDNA_full, the *SMN2* minigene was replaced by a cDNA sequence containing exon 7. In pSMN2_cDNA_Δ7, the *SMN2* minigene was replaced by a cDNA sequence missing exon 7. In plasmids pSMN2_PT_mut1, pSMN2_PT_mut2, and pSMN2_PT_mut3, the purine tract (PT) of exon 7 was disrupted by the respective mutations ex7:G25T,G26T, ex7:Δ22..27, and ex7:Δ17..28. In pSMN2_5ss_consN, the 5'ss of exon 7 was replaced with a consensus 5'ss (NCAG/GUAAGU) having nucleotide N (N=A,C,G, U) at position -4. In pSMN2_5ss_nullN, the 5'ss of exon 7 was replaced with a null 5'ss (NCAG/GGAAGU) having nucleotide N (A,C,G, or U) at position -4. In pSMN1_5ss_mutX, an ex7:T6C mutation was introduced together with an additional mutation X (X=A3C, A3G, A3U, A4C, A4G, A4U, G5A, G5C, G5U) in the intronic region of the exon 7 5'ss.

Minigene plasmid pELP1_WT, which was reported in ref.[1], contains an *ELP1* minigene spanning exons 19, 20, and 21. pELP1_WT also contains the in20:T6C mutation, which reduces exon 20 inclusion and thereby causes familial dysautonomia in affected individuals. In pELP1_cDNA_full, the *ELP1* minigene is replaced by a cDNA sequence containing exon 20. In pELP1_cDNA_Δ20, the *ELP1* minigene is replaced by a cDNA sequence missing exon 20.

##### 1.4 MPSA library construction

The *SMN2* minigene library comprises three sub-libraries (lib1, lib2, lib3), all of which were derived from pSMN2_WT. Each sub-library contains *SMN2* minigenes having 285 variant 5'ss sequences: the wild-type 5'ss, AGGA/GUAAGU; all 24 single-position mutants of the wild-type 5'ss (but with $G_{+1}U_{+2}$ fixed); all 252 two-position mutants of the wild-type 5'ss (but with $G_{+1}U_{+2}$ fixed); 4 consensus 5'ss of the form NCAG/GUAAGU; and 4 null 5'ss of the form NCAG/GGAAGA. The sub-libraries are listed in **Table S2** as pSMN2_5ss_libX (where X=1, 2, 3). **Fig. S1G** shows the resulting number of barcodes associated with each variant 5'ss in each *SMN2* sub-library.

The *ELP1* minigene library comprises two sub-libraries (lib1, lib2), both of which were derived from pELP1_WT. Each sub-library contains *ELP1* minigenes having approximately all 32,768 variant 5'ss sequences of the form ANNN/GYNNNN, where Y indicates C or U. The library was previously reported in ref.[1], which refers to the gene *ELP1* by its old name, *IKBKAP*. The *ELP1* sub-libraries are listed in **Table S2** as pELP1_5ss_libX (where X=1, 2). Each sub-library was independently cloned and therefore has different barcodes associated with each variant 5'ss. **Fig. S2E** shows the resulting number of barcodes associated with each variant 5'ss in each *ELP1* sub-library.

Each sub-library was independently cloned and therefore has different barcodes associated with each variant 5'ss. The sub-libraries were constructed as follows (see also ref.[1])

Cloning of ssbc libraries. Plasmid libraries containing splice-site/barcode (ssbc) fragments were cloned as follows. Equal molar ratios of the oligo pool (*SMN2* ssbc library: SMN2 BseRI ss top, synthesized by IDT; *ELP1* library: ELP1 ssbc BseRI ss top, synthesized by IDT) and a a DNA oligo containing a 20-nt random barcode sequence at the 5' end (*SMN2* ssbc library: SMN2 NotI bc bot; *ELP1* ssbc library: ELP1 XhoI bc bot) were annealed in 1x annealing buffer (10 mM Tris pH 8.0, 50 mM NaCl, 1 mM EDTA) with Phusion High-Fidelity DNA Polymerase (New England Biolabs), heated to 95 ˚C for 5 min, and extended at 72 ˚C for 30 min. The resulting ssbc fragment was then digested (*SMN2* ssbc library: using BseRI and NotI; *ELP1* ssbc library: using BseRI and XhoI) and ligated to a backbone that was amplified by PCR (*SMN2* ssbc library: from pSMN2_WT, using primers SMN2 BB fwd and SMN2 BB rev; *ELP1* ssbc library: from pELP1_WT, using primers ELP1 BB fwd and ELP1 BB rev), digested with DpnI, then digested by the same restriction enzymes. Ligated DNA was purified by drop dialysis using a 0.025 mm membrane filter (Millipore) for at least 2 hr, and electroporated into MegaX DH10B T1 Electrocomp Cells (ThermoFisher) using a 0.1-cm cuvette at 2.0 kV, 200 Ω,25 µF in a BioRad Gene Pulser.

Sequencing of ssbc fragments. To extract ssbc fragments for Illumina sequencing, ssbc libraries were digested (*SMN2* ssbc library: using BsiHKAI and NotI; *ELP1* ssbc library: using PacI and XhoI), and the resulting inserts were ligated to double-stranded DNA fragments containing Illumina-compatible primer sequences (generated by annealing PE1 top with PE1 bot, and PE2 top with PE2 bot). Ligation products were separated on a 2% agarose gel, purified using the QIAquick Gel Extraction Kit (QIAGEN), and sequenced using Illumina HiSeq 2500 PE100 sequencing. The resulting sequence data provided the information needed to associate each 20 nt barcode with a unique 5'ss.

Cloning of minigene libraries. The remaining intronic and exonic sequences were PCR-amplified (*SMN2* minigene library: using primers SMN2 insert F and SMN2 insert R; *ELP1* minigene library: using primers ELP1 insert F and ELP1 insert R), digested with AarI, ligated to AarI-digested ssbc library plasmids.

##### 1.5 MPSA experiments

HeLa cells were treated with 100 nM risdiplam, 50 nM branaplam, or DMSO at the time of transfection. Risdiplam was applied at twice the concentration of branaplam because pilot dose-response experiments revealed that risdiplam is approximately half as potent as branaplam at *SMN2* exon 7. The drug concentrations used in the MPSA experiments correspond to 7.1x the values of $\mathrm{EC}_{2\mathrm{x}}$ (14 nM for risdiplam, 7 nM for branaplam) measured in the pilot dose-response experiments (**Sec. 1.6**, **Fig. S1J,K**).

Each MPSA experiment was carried out as follows. Minigene library DNA (6 µg) was transfected into $5\mathrm{x}10^6$ HeLa cells using Lipofectamine 2000. Transfected cells were then incubated for 48 hr. RNA was then isolated using Trizol. cDNA was synthesized using oligo-dT primer and Improm-II Reverse Transcriptase (Promega). Mulitiple rounds of PCR were then carried out using Phusion High-Fidelity DNA Polymerase as follows. The exon inclusion product was first amplified by PCR using a forward primer that binds the middle exon (*SMN2* minigene library: SMN2 e7F; *ELP1* minigene library: ELP1 e20F) and a minigene-specific reverse primer (BCR). Barcode-containing amplicons were then isolated using a forward primer that binds in the last exon (*SMN2* minigene library: SMN2 e8F; *ELP1* minigene library: ELP1 e21F) and a common reverse primer (barcode R). The same pair of primers were used to directly amplify total RNA barcodes from cDNA. Sample-specific barcodes were then added to exon inclusion barcode products or total RNA barcode product by PCR with the primers barcode-LID F and barcode-LID R. A second round of PCR amplification using primers PE1_v4 and PE2_v4 then added Illumina-compatible ends for sequencing. Both inclusion isoform barcode counts and total RNA barcode counts were obtained via Illumina sequencing, followed by processing using custom Python scripts.

All three *SMN2* sub-libraries were assayed. Three biological replicates (rep1, rep2, rep3) were performed for each sub-library. **Fig. S1H** illustrates the coverage in these MPSA experiments via the number of total isoform reads obtained for each 5’ss in each replicate of each *SMN2* sub-library in each treatment condition.

Both *ELP1* sub-libraries were assayed. Two biological replicates (rep1, rep2) were performed for each sub-library. **Fig. S2G** illustrates the coverage in these MPSA experiments via the number of total isoform reads obtained for each 5’ss in each replicate of each *ELP1* sub-library in each treatment condition.

##### 1.6 Radioactive gel assays

RNA was isolated from minigene-expressing HeLa cells using Trizol. cDNA was made using Improm-II Reverse Transcription System (Promega), following the manufacturer’s instructions. For splicing analysis, a minigene-specific reverse primer (SBCRmod) was used in conjunction with the SMN2 e6F primer in the presence of [$^{32}$P]-dCTP to amplify the splicing isoforms using Q5 High-Fidelity DNA Polymerase (New England Biolabs), following the manufacturer’s instructions. The reaction was initially denatured at 98 ˚C for 2 min, then denatured at 98 ˚C for 15 s, annealed at 60 ˚C for 30 s, and extended at 72 ˚C for 1 min for 20 cycles, with a final extension at 72 ˚C for 10 min. The PCR products were resolved on a 5% non-denaturing polyacrylamide gel and were detected with a Typhoon FLA7000 phosphorimager. Quantification of the isoforms was done using ImageJ (NIH).

**Fig. S1I** shows the radioactive RT-PCR measurements for the four pSMN2_5ss_consN minigenes and the four pSMN2_5ss_nullN minigenes. Data for pSMN2_5ss_consN show that consensus 5’ss give PSI ~100, thereby validating our MPSA normalization method (**Eq. 6** in **Sec. 2.1**). Data for pSMN2_5ss_nullN show that null 5’ss give PSI ~0, thereby verifying that the MPSA has low background.

**Fig. S1J,K** show pilot dose-response experiments for risdiplam and branaplam performed using radioactive RT-PCR. These experiments were used to define the 1x concentrations of each drug for use in designing MPSA, RNA-seq, and drug mixture experiments. Dose-response curves were fit using a Bayesian model analogous to the model in **Sec. 4.4**, but which models PSI instead of inclusion/exclusion ratios. The pilot risdiplam dose-response experiment yielded $\mathrm{EC}_{2\mathrm{x}}$ = 13.5 nM [12.2 nM, 14.9 nM] (**Fig. S1J**); we therefore chose 1x risdiplam = 14 nM. The pilot branaplam dose-response experiment yielded $\mathrm{EC}_{2\mathrm{x}}$ = 7.38 nM [6.35 nM, 8.61 nM] (**Fig. S1K**); we therefore chose 1x branaplam = 7 nM.

##### 1.7 RNA-seq experiments

HeLa cells were treated with 140 nM risdiplam, 70 nM branaplam, or DMSO. These concentrations were chosen to correspond to 10x the $\mathrm{EC}_{2\mathrm{x}}$ values measured for each drug in pilot dose-response experiments. Five biological replicates were performed for each condition. RNA-seq libraries were prepared using KAPA mRNA HyperPrep kits (Roche) and sequenced with one Illumina NextSeq 2000 run using a P3 chip. A total of 1,442,392,333 reads across all 15 RNA samples were obtained.

Illumina sequence data were analyzed using rMATS, which provided estimated PSI values for all 15 samples. We focused our analysis on cassette exon events. Specifically, we selected 247,321 cassette exon events (~68% of the 365,260 events reported by rMATS) that had a median count of at least 10 across all samples and at least a single read supporting exon skipping in any of the samples. These cassette exon events were identified by rMATS de novo—no reference annotation was used to define these events.

To assess the reproducibility of these RNA-seq experiments, we calculated Pearson correlation coefficients for every pair of samples within the same condition. We found Pearson correlation values of $r = 0.988 \pm 0.001$, where $\pm$ indicates standard deviation across pairs of replicates. As this correlation is mainly driven by the large number of exons consistently showing PSI very close to 1 or 0, we also calculated, for each pair of samples, the correlation across exons having 5 < PSI < 95; this yielded $r = 0.889 \pm 0.008$.

Bayesian multiple logistic regression modeling was then used to simultaneously infer PSI values and corresponding 95% posterior credible intervals for all three drug treatments for 235,711 distinct exons having 5’ss sequences with $G_{+1}U_{+1}$. Among these exons were 13,431 distinct 10 nt 5'ss sequences. Other alternative splicing events (including cassette exons with non-GU 5’ss, alternative 5'ss usage, alternative 3'ss usage, and intron retention) were not analyzed.

PSI values were used to infer allelic manifolds describing the effect of each treatment (risdiplam or branaplam) relative to DMSO. Specifically, inference of the Bayesian model described in **Sec. 4.1** was used to infer an allelic manifold for each of the 2,521 5'ss sequences that appeared in at least 10 exons identified by rMATS. The results quantify the estimated effect size $E$, as well as 95% posterior credible intervals for $E$, for each 5'ss sequence under treatment with risdiplam (effect size $E_{\text{ris}}$) or branaplam (effect size $E_{\text{bran}}$).

##### 1.8 qPCR assays

Improm-II™ RT (Promega) was applied to 1 µg DNase-treated total RNA using the manufacturer's protocol and using the minigene-specific primer BCR_short (**Table S3**). The qPCR mixture, at a final volume of 10 µL in a 384-well qPCR plate, consisted of 5 µL PowerUP SYBR Green Master Mix (Thermo Fisher Scientific), 0.2 µL ROX reference dye (Thermo Fisher Scientific), 0.3 µL nuclease-free water, 2 µL primer mix (5 µM forward primer, 5 µM reverse primer), and 2.5 µL of 20 ng/µl total RNA. The qPCR reactions were performed on a QuantStudio™ 6 Flex System (ThermoFisher Scientific) with thermal cycling as follows: 50 ˚C for 2 minutes, 95 ˚C for 10 minutes, 40 cycles of denaturation at 95 ˚C for 15 seconds and extension at 60 ˚C for 1 minute. Three to six technical replicates were carried out for every biological replicate. The resulting $C_t$ values were exported as an Excel spreadsheet for analysis using custom Python scripts (available on GitHub).

qPCR reactions were carried out using isoform-specific forward primers and SBCRmod (**Table S3**) as the reverse primer. For *SMN2* minigenes, inclusion isoforms were quantified using forward primer SMN_inclusion_fwd, while exclusion isoforms were quantified using SMN_exclusion_fwd (**Table S3**). Forward primers were validated via RT-qPCR experiments on cells transiently transfected with plasmids pSMN2_cDNA_full and pSMN2_cDNA_Δ7 (**Table S2**). For *ELP1* minigenes, inclusion isoforms were quantified using forward primer ELP1_inclusion_fwd, while exclusion isoforms were quantified using ELP1_exclusion_fwd (**Table S3**). These primer pairs were validated via RT-qPCR experiments on cells transiently transfected with plasmids pELP1_cDNA_full and pELP1_cDNA_Δ20 (**Table S2**). Standard curves were measured using a dilution series of template plasmid, and confirmed that all qPCR primer pairs had amplification efficiencies that were statistically indistinguishable from 100% on the correct cDNA transcript and negligible on the incorrect cDNA transcript.

For **Fig. 4F,G**, drug effect values $E$ were computed using $E = 2^{-\Delta\Delta C_t}$, where

$$\Delta\Delta C_t = \left(C_t^{\text{inclusion,drug}} - C_t^{\text{exclusion,drug}}\right) - \left(C_t^{\text{inclusion,DMSO}} - C_t^{\text{exclusion,DMSO}}\right), \quad (1)$$

and where the four $C_t$ values were measured for two different isoforms (inclusion and exclusion) in two different conditions (drug and DMSO).

##### 1.9 Dose-response experiments and linear-mixture experiments

Serial dilutions of risdiplam and branaplam were prepared from 1 mM stock solutions in DMSO. Serial dilutions of RECTAS were prepared from 50 mM stock solutions in DMSO. Serial dilutions of ASOi6, ASOi7, and ASOi20 were prepared from 100 µM stock solutions in de-ionized water.

At each assayed drug concentration, the abundances of inclusion and exclusion transcripts in total extracted RNA were quantified using RT-qPCR with isoform-specific primers as described in **Sec. 1.8**. Two biological replicates were assayed at each drug concentration. For each biological replicate, three technical replicates were performed, after which inclusion/exclusion ratios were quantified as

$$\frac{\text{inclusion}}{\text{exclusion}} = 2^{-\Delta \text{C}_\text{t}}, \qquad \Delta C_t = C_t^{\text{inclusion}} - C_t^{\text{exclusion}}, \quad (2)$$

where $C_t^{\text{inclusion}}$ and $C_t^{\text{exclusion}}$ denote the median qPCR results respectively obtained using inclusion-specific or exclusion-specific forward primers. Below we list the drug concentrations assayed for each dose-response curve. Note: 1x risdiplam is 14 nM; 1x branaplam is 7 nM; 1x ASOi6 is 0.6 nM; 1x ASOi7 is 0.1 nM; and 1x cocktail is defined as a mixture comprising 0.5x of each of the two component drugs.

**Fig. 5E-L:**

- Risdiplam dose-response curves (**Fig. 5E-H**) were measured on *SMN2* minigenes using 1,000 nM, 500 nM, 250 nM,125 nM, 50 nM, 25 nM, 12.5 nM, 5 nM, 2.5 nM,1.25 nM, 0.5 nM, 0.25 nM, and 0 nM risdiplam, or a subset of these concentrations.

- Branaplam dose-response curves (**Fig. 5I-L**) were measured on *SMN2* minigenes using 1,000 nM, 500 nM, 250 nM,125 nM, 50 nM, 25 nM, 12.5 nM, 5 nM, 2.5 nM,1.25 nM, 0.5 nM, 0.25 nM, and 0 nM branaplam, or a subset of these concentrations.

**Fig. 6A-D:**

- The ASOi7 dose-response curve (**Fig. 6A**) was measured on the *SMN2* WT minigenes using 100 nM, 50 nM, 25 nM, 12.5 nM, 5 nM, 2.5 nM, 1.25 nM, 0.5 nM, 0.25 nM, 0.125 nM, 0.05 nM, 0.025 nM, 0.0125 nM, and 0 nM ASOi7.
- The ASOi6 dose-response curve (**Fig. 6B**) was measured on the *SMN2* WT minigenes using 100 nM, 50 nM, 25 nM, 12.5 nM, 5 nM, 2.5 nM, 1.25 nM, 0.5 nM, 0.25 nM, 0.125 nM, 0.05 nM, and 0 nM ASOi6.
- The RECTAS dose-response curve (**Fig. 6C**) was measured on the *ELP1* WT minigene using 12,500 nM, 5,000 nM, 2,500 nM, 1,250nM, 500 nM, 250 nM, 125 nM, 50 nM, and 0 nM RECTAS.
- The ASOi20 dose response curve (**Fig. 6D**) was measured on the *ELP1* WT minigene using 10 nM, 5 nM, 2.5 nM, 1 nM, 0.5 nM, 0.25 nM, 0.1 nM, 0.05 nM, 0.025 nM, 0 nM.

**Fig. 6E-K:**

- The risdiplam/branaplam linear ramp curve (**Fig. 6E**) was measured on the *SMN2* WT minigene using drug concentrations of 0x/10x, 2x/8x, 4x/6x, 5x/5x, 6x/4x, 8x/2x, and 10x/0x.
- The risdiplam/ASOi6 linear ramp curve (**Fig. 6F**) was measured on the *SMN2* WT minigene using drug concentrations of 0x/50x, 2x/40x, 4x/30x, 5x/25x, 6x/20x, 8x/10x, and 10x/0x.
- The branaplam/ASOi6 linear ramp curve (**Fig. 6G**) was measured on the *SMN2* WT minigene using drug concentrations of 0x/50x, 2x/40x, 4x/30x, 5x/25x, 6x/20x, 8x/10x, and 10x/0x.
- The risdiplam/ASOi7 linear ramp curve (**Fig. 6H**) was measured on the *SMN2* WT minigene using drug concentrations of 0x/100x, 5x/80x, 10x/60x, 12.5x/50x, 15x/40x, 20x/20x, and 25x/0x.
- The branaplam/ASOi7 linear ramp curve (**Fig. 6I**) was measured on the *SMN2* WT minigene using drug concentrations of 0x/100x, 10x/80x, 20x/60x, 25x/50x, 30x/40x, 40x/20x, and 50x/0x.
- The ASOi6/ASOi7 linear ramp curve (**Fig. 6J**) was measured using on the *SMN2* WT minigene drug concentrations of 0x/25x, 10x/20x, 20x/15x, 25x/12.5x, 30x/10x, 40x/5x, and 50x/0x.
- The RECTAS/ASOi20 linear ramp curve (**Fig. 6K**) was measured on the *ELP1* WT minigene using drug concentrations of 0x/25x, 5x/20x, 10x/15x, 12.5x/12.5x, 15x/10x, 20x/5x, and 25x/0x.

**Fig. S12A-F:**

- The negative-control risdiplam dose-response curve (**Fig. S12A**) was measured on the *ELP1* WT minigene using drug concentrations of 1,000 nM, 100 nM, 10 nM, and 0 nM.
- The negative-control branaplam dose-response curve (**Fig. S12B**) was measured on the *ELP1* WT minigene using drug concentrations of 500 nM, 50 nM, 5 nM, and 0 nM.
- The negative-control ASOi6 dose-response curve (**Fig. S12C**) was measured on the *ELP1* WT minigene using drug concentrations of 50 nM, 5 nM, 0.5 nM, and 0 nM.
- The negative-control ASOi7 dose-response curve (**Fig. S12D**) was measured on the *ELP1* WT minigene using drug concentrations of 50 nM, 5 nM, 0.5 nM, and 0 nM.
- The negative-control RECTAS dose-response curve (**Fig. S12E**) was measured on the *SMN2* WT minigene using drug concentrations of 25 μM, 2.5 μM, 0.25 μM, and 0 μM.
- The negative-control ASOi20 dose-response curve (**Fig. S12F**) was measured on the *SMN2* WT minigene using drug concentrations of 10 nM, 1 nM, 0.1 nM, and 0 nM.

**Fig. S13A-G:**

- The risdiplam/branaplam cocktail dose-response curve (**Fig. S13A**) was measured on the *SMN2* WT exon 7 minigene using cocktail concentrations of 100x, 50x, 25x, 10x, 5x, 2.5x, 1x, 0.5x, and 0x.

- The risdiplam/ASOi6 cocktail dose-response curve (**Fig. S13B**) was measured on the *SMN2* WT minigene using cocktail concentrations of 250x, 100x, 50x, 25x, 10x, 5x, 2.5x, 1x, 0.5x and 0x.
- The risdiplam/ASOi7 cocktail dose-response curve (**Fig. S13C**) was measured on the *SMN2* WT minigene using cocktail concentrations of 250x, 100x, 50x, 25x, 10x, 5x, 2.5x, 1x, 0.5x and 0x.
- The branaplam/ASOi6 cocktail dose-response curve (**Fig. S13D**) was measured on the *SMN2* WT minigene using cocktail concentrations of 250x, 100x, 50x, 25x, 10x, 5x, 2.5x, 1x, 0.5x, and 0x.
- The branaplam/ASOi7 cocktail dose-response curve (**Fig. S13E**) was measured on the *SMN2* WT minigene using cocktail concentrations of 250x, 100x, 50x, 25x, 10x, 5x, 2.5x, 1x, 0.5x, and 0x.
- The ASOi6/ASOi7 cocktail dose-response curve (**Fig. S13F**) was measured on the *SMN2* WT minigene using cocktail concentrations of 250x, 100x, 50x, 25x, 10x, 5x, 2.5x, 1x, and 0x.
- The RECTAS/ASOi20 cocktail dose-response curve (**Fig. S13G**) was measured on the *ELP1* WT minigene using cocktail concentrations of 20x, 10x, 5x, 2.5x, 1x, 0.5x, 0.2x, and 0x.

#### *2 Data processing, exploratory analysis, and molecular dynamics simulations*

##### 2.1 Computation of PSI values from MPSA data

In what follows, $l$ indexes the three assayed *SMN2* minigene sub-libraries (lib1, lib2, lib3), $r$ indexes the three biological replicate experiments (rep1, rep2, rep3) performed for each sub-library, $s$ indexes the 285 assayed 5'ss sequences in the *SMN2* library, and $b$ indexes 20 nt random barcode sequences. $n_{s,b}^{\text{ssbc},l}$ denotes the number of splice-site-barcode (ssbc) reads obtained for sub-library $l$ that have 5'ss $s$ and barcode $b$. $n_b^{\text{tot},l,r}$ denotes the number of total (tot) mRNA reads obtained for replicate $r$ of the experiment on sub-library $l$ that contain barcode $b$. $n_b^{\text{inc},l,r}$ denotes the number of exon inclusion (inc) mRNA reads, obtained for replicate $r$ of the experiment on sub-library $l$ that contain barcode $b$. $s_l(b)$ denotes the 5'ss sequence associated with barcode $b$ in sub-library $l$.

We associated barcodes with 5'ss as follows. In each sub-library $l$, barcode $b$ was associated with 5'ss $s$ if and only if

$$n_{s,b}^{\text{ssbc},l} \geq 2 \quad \text{and} \quad n_{s,b}^{\text{ssbc},l} \geq 4 \times \sum_{s' \neq s} n_{s',b}^{\text{ssbc},l}, \tag{3}$$

i.e., barcode $b$ was linked to 5'ss $s$ in at least 2 reads and in at least 4 times as many reads as barcode $b$ was linked to any other 5'ss $s'$. When these conditions were met, we defined $s_l(b) \equiv s$.

We computed PSI values as follows. For each sub-library $l$, each replicate $r$, and each 5'ss $s$, we computed the number of total mRNA reads $n_s^{\text{tot},l,r}$, and the number of exon inclusion mRNA reads $n_s^{\text{inc},l,r}$, that contained barcodes associated with $s$ using

$$n_s^{\text{tot},l,r} = \sum_{\{b:s_{l(b)}=s\}} n_b^{\text{tot},l,r} \quad \text{and} \quad n_s^{\text{inc},l,r} = \sum_{\{b:s_{l(b)}=s\}} n_b^{\text{inc},l,r}. \tag{4}$$

For values of $s$, $l$, and $r$ such that $n_s^{\text{tot},l,r} \geq 5$, we then computed the read ratio

$$\rho_s^{l,r} = \frac{n_s^{\text{inc},l,r}}{n_s^{\text{tot},l,r}}. \tag{5}$$

We then compute the corresponding PSI value as

$$\Psi_s^{l,r} = 100 \times \frac{\rho_s^{l,r}}{\rho_{\text{cons}}^{l,r}}, \tag{6}$$

where $\rho_{\text{cons}}^{l,r}$ is the mean read ratio $\rho_s^{l,r}$ for the four consensus 5'ss sequences present in the *SMN2* library. For each 5'ss $s$, we then estimated a single PSI value by computing

$$\Psi_s = 100 \times \frac{\widetilde{\Psi}_s}{\widetilde{\Psi}_{\text{cons}}} \quad \text{where} \quad \widetilde{\Psi}_s = \text{median}_{\{l,r\}}\left(\Psi_s^{l,r}\right). \tag{7}$$

where $\widetilde{\Psi}_{\text{cons}}$ is the median $\widetilde{\Psi}_s$ value over the four consensus sequences. We also report a standard error in log PSI,

$$\sigma_s = \text{SE}_{\{l,r\}}(\log \Psi_s^{l,r}), \tag{8}$$

where SE is computed over all $l, r$ paris for which $\Psi_s^{l,r}$ is above a minimum threshold of $\Psi_{\min} = 10^{-2}$. The standard errors were used to compute the uncertainties shown in **Fig. S1A-F**.

An analogous method was used to compute PSI values for the *ELP1* MPSA data, with the following changes: the data comprised two replicates (rep1, rep2) for each of two sub-libraries (lib1, lib2), a threshold of $n_s^{\text{tot},l,r} \geq 2$ was used to compute read ratios, and $\rho_{\text{cons}}^{l,r}$ was set equal to the read ratio $\rho_s^{l,r}$ where $s$ denotes 5'ss sequence ACAA/GUAAGU.

#### 2.2 Classification of 5'ss from MPSA data

Each variant of the *SMN2* exon 7 5'ss was classified based on PSI values measured by MPSA on cells treated with risdiplam, branaplam, or DMSO. In what follows, these PSI values are denoted by $PSI_{ris}$, $PSI_{bran}$, and $PSI_{DMSO}$. Each class of 5'ss was defined in terms of thresholds on individual PSI values, together with thresholds on ratios of PSI values after subtracting background. Background PSI values were computed as the median of the PSI values measured for the four null 5'ss; these background PSI values were $PSI_{ris,bg}$=0.27, $PSI_{bran,bg}$=0.38, and $PSI_{DMSO,bg}$=0.30.

In **Fig. 1D** and **Figs. S3A,B**, the following thresholds were used:

- Class 1-ris: $0.67 < PSI_{ris} < 67$, $0.67 < PSI_{DMSO} < 67$, and $(PSI_{ris} - PSI_{ris,bg})/ (PSI_{DMSO}-PSI_{DMSO,bg}) > 4$.
- Class 2-ris: $0.67 < PSI_{ris} < 67$, $0.67 < PSI_{DMSO} < 67$, and $1/3 < (PSI_{ris} - PSI_{ris,bg})/ (PSI_{DMSO}-PSI_{DMSO,bg}) < 3$.

In **Fig. 1E** and **Figs. S3C,D**, the following thresholds were used:

- Class 1-hyp: $1.2 < PSI_{bran} < 40$; $1.2 < PSI_{ris} < 40$; $(PSI_{bran} - PSI_{bran,bg})/ (PSI_{ris}-PSI_{ris,bg}) > 6$.
- Class 2-hyp: $1.2 < PSI_{bran} < 40$; $1.2 < PSI_{ris} < 40$; $1/3 < (PSI_{bran} - PSI_{bran,bg})/ (PSI_{ris}-PSI_{ris,bg}) < 3$.

In **Fig. 1F**, and **Figs. S3E**,**F**, the following thresholds were used:

- Class 1-bran: $1.2 < PSI_{bran} < 40$; $1.2 < PSI_{DMSO} < 40$; $(PSI_{bran} - PSI_{bran,bg})/ (PSI_{DMSO}-PSI_{DMSO,bg}) > 6$.
- Class 2-bran: $1.2 < PSI_{bran} < 40$; $1.2 < PSI_{DMSO} < 40$; $1/3 < (PSI_{bran} - PSI_{bran,bg})/ (PSI_{DMSO}-PSI_{DMSO,bg}) < 3$.

In **Figs. S4A-C**, we used the same minimum and maximum PSI thresholds as in **Fig. 1F**, but varied the thresholds placed on the ratios of background-subtracted PSI values as follows:

- **Fig. S4A**:
  - Class 1-bran: $(PSI_{bran} - PSI_{bran,bg})/ (PSI_{DMSO}-PSI_{DMSO,bg}) > 4$.
  - Class 2-bran: $1/3 < (PSI_{bran} - PSI_{bran,bg})/ (PSI_{DMSO}-PSI_{DMSO,bg}) < 3$.
- **Fig. S4B**:
  - Class 1-bran: $(PSI_{bran} - PSI_{bran,bg})/ (PSI_{DMSO}-PSI_{DMSO,bg}) > 6$.
  - Class 2-bran: $1/2 < (PSI_{bran} - PSI_{bran,bg})/ (PSI_{DMSO}-PSI_{DMSO,bg}) < 2$.
- **Fig. S4C**:
  - Class 1-bran: $(PSI_{bran} - PSI_{bran,bg})/ (PSI_{DMSO}-PSI_{DMSO,bg}) > 8$.
  - Class 2-bran: $1/1.5 < (PSI_{bran} - PSI_{bran,bg})/ (PSI_{DMSO}-PSI_{DMSO,bg}) < 1.5$.

#### 2.3 Inference of restrictive and permissive IUPAC motifs from MPSA data

Both the risdiplam classification criteria and hyper-activation classification criteria were satisfied by multiple IUPAC motifs. To characterize the range of IUPAC motifs consistent with each classification criterion, we further determined a "restrictive" IUPAC motif and "permissive" IUPAC motif: the restrictive IUPAC motif matches the fewest 5'ss sequences while being consistent with a given classification criterion; the permissive IUPAC motif matches the most 5'ss sequences consistent with a given classification criterion. Formally, the restrictive and permissive IUPAC motifs were defined as follows.

The restrictive IUPAC motif was defined as the IUPAC motif having $G_{+1}U_{+2}$ that (i) matches all 5'ss in class 1, (ii) matches no 5'ss in class 2, and (iii) matches as few 5'ss sequences as possible. This motif was computationally determined using a simple deterministic algorithm: for each 5'ss nucleotide position $l$ from -4 to +6 (except +1 and +2), the set of bases allowed by the motif at position $l$ was set equal to the set bases at position $l$ that appear in 5'ss in class 1. It is readily seen that, at each position $l$, the set of nucleotides permitted by the resulting motif cannot be further restricted without excluding at least one 5'ss in class 1, thus proving that this procedure identifies a unique motif. We then verified that the resulting motif does not match any 5'ss in class 2.

The permissive IUPAC motif was defined as the IUPAC motif having $G_{+1}U_{+2}$ that (i) matches all 5'ss in class 1, (ii) matches no 5'ss in class 2, and (iii) matches as many 5'ss sequences in class 3 as possible. This motif was computationally determined using the following non-deterministic greedy algorithm. Starting from the corresponding restrictive IUPAC motif, the algorithm randomly chose a 5'ss nucleotide position $l$ from -4 to +6 (except +1 and +2) and base $b$ such that base $b$ is not permitted by the motif at position $l$. The motif was then augmented to permit base $b$ at position $l$, and this augmented motif was kept if it still satisfied criteria i and ii. This loop iterated until it was not possible to augment the motif any further without violating criteria i or ii. We reran this greedy algorithm 100 times. In each application, all 100 runs of the greedy algorithm yielded the same permissive risdiplam IUPAC motif (**Fig. S3B**) or permissive hyper-activation IUPAC motif (**Fig. S3D**).

##### 2.4 Molecular dynamics simulations

The molecular dynamics simulations featured in **Fig. S10** were carried out as follows. Risdiplam (**Fig. S10A**) and the enol tautomer of branaplam (**Fig. S10B**) were parametrized with OpenFF[2]. One microsecond of simulation was conducted for each drug molecule in a box of water with 150 mM of NaCl. All simulations were set up using CHARMM-GUI[3]. Simulations were then performed with the GROMACS 2022 software package[4]. The Charmm36m[5] forcefield with the TIP3P water model was used. Each system was energy minimized using steepest descent followed by a multi-step equilibration lasting a total of 375 ps. In the first two steps, the system was equilibrated in a canonical (NVT) ensemble with an integration time step of 1 fs for 50 ps each, maintaining a temperature of 303.15 K using the Berendsen thermostat[6]. During the equilibration steps, we established a pressure of 1 bar using the Berendsen barostat[6] with a characteristic time of 1 ps. During the production simulations, no restraints were applied on the systems. We applied the velocity rescale thermostat[7] with a characteristic time of 1 ps to keep a constant temperature of 303.15 K. We applied the Parrinello-Rahman barostat[8] and a characteristic time of 5 ps to keep a constant pressure of 1 bar. Simulations were analyzed in Python using MDTraj[9].

#### 3 *Mathematical model definitions*

Here we mathematically define the four biophysical models presented in **Fig. 2** and **Fig. S5** (**Sec. 3.1**), **Fig. 3** (**Sec. 3.2**), **Fig. S6** (**Sec. 3.3**), and **Fig. S11** (**Sec. 3.4**). These models assume that PSI is equal to 100 times the occupancy of the 5'ss by U1 snRNP in thermodynamic equilibrium. In what follows, we let $x$ denote the 10 nt sequence of a 5'ss, $z$ denote the surrounding pre-mRNA sequence (i.e., "context"), and $y$ denote the identity and concentration of the applied drug.

##### 3.1 Biophysical allelic-manifold model

The biophysical allelic-manifold model, illustrated in **Fig. 2** and **Fig. S5**, assumes that pre-mRNA containing an exon of interest can be in one of three states.

- State 1 corresponds to pre-mRNA not bound by U1 at the 5'ss; this state is assigned Gibbs free energy of 0.
- State 2 corresponds to exon pre-mRNA bound by U1 at the 5'ss; this state is assigned a Gibbs free energy of $\Delta G_{\text{U1}}(x,z)$, which depends on the sequence of the 5'ss, $x$, and on the surrounding pre-mRNA sequence context, $z$ (e.g., due to the effects of splicing enhancers, splicing silencers, etc.), but not on the identity and concentration of the drug, $y$.
- State 3 corresponds to exon pre-mRNA bound by U1 at the 5'ss and bound by drug; this state is assigned a Gibbs free energy of $\Delta G_{\text{U1}}(x,z) + \Delta G_{\text{drug}}(x,y)$, where $\Delta G_{\text{drug}}(x,y)$ depends on the sequence of the 5'ss, $x$, and the identity and concentration of the drug, $y$, but not on the surrounding pre-mRNA sequence context, $z$.

PSI is then given by 100 times the fractional occupancy of U1 in thermal equilibrium, i.e.

$$\Psi(x,y,z) = 100 \times \frac{e^{-\Delta G_{\text{U1}}(x,z)/RT} + e^{-[\Delta G_{\text{U1}}(x,z)+\Delta G_{\text{drug}}(x,y)]/RT}}{1 + e^{-\Delta G_{\text{U1}}(x,z)/RT} + e^{-[\Delta G_{\text{U1}}(x,z)+\Delta G_{\text{drug}}(x,y)]/RT}}, \tag{9}$$

where $R = 1.987 \times 10^{-3} \frac{\text{kcal}}{\text{mol·K}}$ is the gas constant and $T = 310$ K is temperature (and thus $RT = 0.616 \frac{\text{kcal}}{\text{mol}}$). Defining "context strength" $S$ and "drug effect" $E$ as

$$S(x,z) = e^{-\Delta G_{\text{U1}}(x,z)/RT}, \qquad E(x,y) = 1 + e^{-\Delta G_{\text{drug}}(x,y)/RT}, \tag{10}$$

we get

$$\Psi(x,y,z) = 100 \times \frac{S(x,z)E(x,y)}{1 + S(x,z)E(x,y)}. \tag{11}$$

Note that, when no drug is present, $\Delta G_{\text{drug}} = -\infty$ and $E = 1$. We thus recover the equations in **Fig. 3** from those in **Fig. S5.**

The allelic manifolds illustrated in **Fig. 3** are formulated as follows. Our RNA-seq experiments measured PSI in three conditions: no drug (condition $y_{\text{DMSO}}$), 140 nM risdiplam (condition $y_{\text{ris}}$), and 70 nM branaplam (condition $y_{\text{bran}}$). Given a fixed 5'ss sequence, $x$, one can write

$$\Psi_{\text{DMSO}}(S) = 100 \times \frac{S}{1+S}, \qquad \Psi_{\text{ris}}(S) = 100 \times \frac{SE_{\text{ris}}}{1+SE_{\text{ris}}}, \qquad \Psi_{\text{bran}}(S) = 100 \times \frac{SE_{\text{bran}}}{1+SE_{\text{bran}}} \tag{12}$$

where $E_{\text{ris}} = E(x, y_{\text{ris}})$, $E_{\text{bran}} = E(\text{x}, \text{y}_{\text{bran}})$, and $S = S(x,z)$. Letting $S$ take on all possible positive values, these equations define a one-dimensional curve (an "allelic manifold", see ref.[10]) in the three-dimensional space defined by the coordinates $(\Psi_{\text{DMSO}}, \Psi_{\text{ris}}, \Psi_{\text{bran}})$.

The allelic manifold formulation is useful because it reveals that, if the biophysical model is correct, then measurements plotted in the three-dimensional space having coordinates $(\Psi_{\text{DMSO}}, \Psi_{\text{ris}}, \Psi_{\text{bran}})$ should collapse to a one-dimensional curve. The shape of the allelic manifold is determined by the drug effect values $E_{\text{ris}}$ and $E_{\text{bran}}$, which are determined by 5'ss sequence $x$ and drug treatment $y$. By contrast, locations along the length of the allelic manifold are determined by context strength $S$, which is determined by context $z$. The term "allelic" refers to the different sequence contexts $z$ acting as an allelic series. The allelic manifold formulation thus separates the influence of context strength from 5'ss strength and drug treatment. The coordinates

($\Psi_{\text{DMSO}}, \Psi_{\text{ris}}, \Psi_{\text{bran}}$) act as different phenotypes for alleles in that series. Note that, if $\Psi_{\text{DMSO}}$, $\Psi_{\text{ris}}$, and $\Psi_{\text{bran}}$ are measured for a given 5'ss $x$ in $N > 1$ different genomic contexts $z$, then **Eq. 11** provides $3N$ nonlinear constraints on $N + 2$ parameters, thereby allowing the determination of $E_{\text{ris}}$, $E_{\text{bran}}$ and all $S$ values. This system of nonlinear equations is what enables the Bayesian inference procedure (described in **Sec. 4.1**) to determine the values for $E_{\text{ris}}$ and $E_{\text{bran}}$ that are plotted in **Fig. 2D**.

The biophysical allelic manifold model thus has 194,129 parameters: context strength values $S$ for 189,087 different genomic exons, and drug effect values $E_{\text{ris}}$ and $E_{\text{bran}}$ for 2,521 distinct 5'ss sequences. This model also includes 9 hyperparameters describing experimental noise; see **Sec. 4.1** for details.

##### 3.2 Biophysical two-interaction-mode model

The two-interaction-mode model for drug effect, illustrated in **Fig. 3**, is a thermodynamic model that explicitly describes drug effect in terms of 5'ss sequence. The name comes from the fact that the model assumes branaplam interacts with the U1/5'ss complex in two distinct interaction modes. The pre-mRNA states and corresponding Gibbs free energies assumed by the model are as follows:

- In the presence of DMSO, two states are possible: 5'ss not bound by U1 (state 1, Gibbs free energy 0) and 5'ss bound by U1 but not by risdiplam (state 2, Gibbs free energy $\Delta G_{\text{U1}}(x,z)$).
- In the presence of risdiplam, three states are possible: state 1, state 2, and 5'ss bound by U1 and by risdiplam (state 3, Gibbs free energy $\Delta G_{\text{U1}}(x,z) + \Delta G_{\text{ris}}(x)$).
- In the presence of branaplam, four states are possible: state 1, state 2, 5'ss bound by U1 and by branaplam in the risdiplam interaction mode (state 4, Gibbs free energy $\Delta G_{\text{U1}}(x,z) + \Delta G_{\text{ris}}(x)$), and 5'ss bound by U1 and by branaplam in the hyper-activation interaction mode (state 5, Gibbs free energy $\Delta G_{\text{U1}}(x,z) + \Delta G_{\text{hyp}}(x)$)

The Gibbs free energies are parameterized as follows. $\Delta G_{\text{U1}}(x,z)$ is assumed to be independent for every combination of 5'ss sequence $x$ and context $z$. Formally,

$$\Delta G_{\text{U1}}(x,z) = \theta^{\text{U1}}_{xz} \tag{13}$$

for some parameter $\theta^{\text{U1}}_{xz}$. $\Delta G_{\text{ris}}(x)$ is assumed to depend on the sequence of the 5'ss via a "risdiplam energy motif". Formally,

$$\Delta G_{\text{ris}}(x) = \theta_0^{\text{ris}} + \sum_{l,b} \theta_{l:b}^{\text{ris}}\, x_{l:b}, \tag{14}$$

where $l$ indexes nucleotide positions within the 5'ss, $b$ indexes the four possible RNA bases ($\text{A, C, G, U}$), $\theta_0^{\text{ris}}$ and $\theta_{l:b}^{\text{ris}}$ are the constant and additive parameters of the risdiplam energy motif, and $x_{l:b}$ is the one-hot encoding of 5'ss sequence (equal to 1 if base $b$ occurs at position $l$, equal to 0 otherwise). Similarly, $\Delta G_{\text{hyp}}(x)$ is assumed to depend on the sequence of the 5'ss via a "hyper-activation energy motif". Formally,

$$\Delta G_{\text{hyp}}(x) = \theta_0^{\text{hyp}} + \sum_{l,b} \theta_{l:b}^{\text{hyp}}\, x_{l:b}, \tag{15}$$

where $\theta_0^{\text{hyp}}$ and $\theta_{l:b}^{\text{hyp}}$ are the constant and additive parameters of the hyper-activation energy motif. A hard-coded parameter, $\Delta\mu = RT \log\,(10/7.1) = 0.211\ \text{kcal/mol}$, accounts for the fact that 10x drug concentrations were applied in the RNA-seq experiment, whereas 7.1x drug concentrations were applied in the MPSA experiments.

Experimental measurements are predicted in terms of Gibbs free energies as follows. Drug effect of 10x risdiplam, as measured by RNA-seq followed by allelic manifold inference, is predicted using

$$E_{\text{ris}} = 1 + e^{[-\Delta G_{\text{ris}}(x) + \Delta\mu]/RT}. \tag{16}$$

Drug effect of 10x branaplam, as measured by RNA-seq followed by allelic manifold inference, is predicted using

$$E_{\text{bran}} = 1 + e^{[-\Delta G_{\text{ris}}(x) + \Delta\mu]/RT} + e^{[-\Delta G_{\text{hyp}}(x) + \Delta\mu]/RT}. \tag{17}$$

PSI in the absence of drug, as measured by MPSA, is predicted using

$$\Psi_{\text{DMSO}} = 100 \times \frac{e^{-\Delta G_{\text{U1}}(x,z)/RT}}{1 + e^{-\Delta G_{\text{U1}}(x,z)/RT}}. \tag{18}$$

PSI in the presence of 7.1x risdiplam, as measured by MPSA, is predicted using

$$\Psi_{\text{ris}} = 100 \times \frac{e^{-\Delta G_{\text{U1}}(x,z)/RT} + e^{-[\Delta G_{\text{U1}}(x,z)+\Delta G_{\text{ris}}(x)]/RT}}{1 + e^{-\Delta G_{\text{U1}}(x,z)/RT} + e^{-[\Delta G_{\text{U1}}(x,z)+\Delta G_{\text{ris}}(x)]/RT}}. \tag{19}$$

PSI in the presence of 7.1x branaplam, as measured by MPSA, is predicted using

$$\Psi_{\text{bran}} = 100 \times \frac{e^{-\Delta G_{\text{U1}}(x,z)/RT} + e^{-[\Delta G_{\text{U1}}(x,z)+\Delta G_{\text{ris}}(x)]/RT} + e^{-[\Delta G_{\text{U1}}(x,z)+\Delta G_{\text{hyp}}(x)]/RT}}{1 + e^{-\Delta G_{\text{U1}}(x,z)/RT} + e^{-[\Delta G_{\text{U1}}(x,z)+\Delta G_{\text{ris}}(x)]/RT} + e^{-[\Delta G_{\text{U1}}(x,z)+\Delta G_{\text{hyp}}(x)]/RT}}. \tag{20}$$

We note that this model is consistent with the allelic manifold model illustrated in **Fig. 2** and **Fig. S5**. $\Delta G_{\text{U1}}$ is the same in both models, and the relevant values of $\Delta G_{\text{drug}}$ are given in terms of $\Delta G_{\text{ris}}$ and $\Delta G_{\text{bran}}$ by

$$\Delta G_{\text{drug}}(x, y_{\text{ris}}) = \Delta G_{\text{ris}}(x), \qquad \Delta G_{\text{drug}}(x, y_{\text{bran}}) = -RT \log \left[e^{-\Delta G_{\text{ris}}(x)/RT} + e^{-\Delta G_{\text{bran}}(x)/RT}\right], \tag{21}$$

where $y_{\text{ris}}$ denotes the presence of 7.1x risdiplam and $y_{\text{bran}}$ denotes the presence of 7.1x branaplam.

The two-interaction-mode model for drug effect thus has 351 parameters that were inferred from MPSA and RNA-seq data: 285 parameters $\theta^{\text{U1}}_{xz}$ (one context $z$, representing the *SMN2* minigene, and 285 variant 5'ss sequences $x$), 32 additive parameters $\theta^{\text{ris}}_{l:b}$, 32 additive parameters $\theta^{\text{hyp}}_{l:b}$, 1 constant parameter $\theta^{\text{ris}}_0$, and 1 constant parameter $\theta^{\text{hyp}}_0$. The model also have 5 hyperparameters describing experimental noise; see **Sec. 4.2** for details.

##### 3.3 Biophysical one-interaction-mode model

The one-interaction-mode model for drug effect, illustrated in **Fig. S6**, is a thermodynamic model that explicitly describes drug effect in terms of 5'ss sequence. The name comes from the fact that the model assumes branaplam interacts with the U1/5'ss complex in only one interaction mode (as opposed to two). The pre-mRNA states and corresponding Gibbs free energies assumed by the model are as follows:

- In the presence of DMSO, two states are possible: 5'ss not bound by U1 (state 1, Gibbs free energy 0) and 5'ss bound by U1 but not by risdiplam (state 2, Gibbs free energy $\Delta G_{\text{U1}}(x,z)$).
- In the presence of risdiplam, three states are possible: state 1, state 2, and 5'ss bound by U1 and by risdiplam (state 3, Gibbs free energy $\Delta G_{\text{U1}}(x,z) + \Delta G_{\text{ris}}(x)$).
- In the presence of branaplam, three states are possible: state 1, state 2, and 5'ss bound by U1 and by branaplam (state 4, Gibbs free energy $\Delta G_{\text{U1}}(x,z) + \Delta G_{\text{bran}}(x)$).

The Gibbs free energies are parameterized as follows. $\Delta G_{\text{U1}}(x,z)$ and $\Delta G_{\text{ris}}(x)$ are defined as in the two-interaction-mode model (**Sec. 3.2**). $\Delta G_{\text{bran}}(x)$ is assumed to depend on the sequence of the 5'ss via a branaplam energy motif. Formally,

$$\Delta G_{\text{bran}}(x) = \theta^{\text{bran}}_0 + \sum_{l,b} \theta^{\text{bran}}_{l:b}\, x_{l:b}, \tag{22}$$

where $\theta^{\text{bran}}_0$ and $\theta^{\text{bran}}_{l:b}$ are the constant and additive parameters of the branaplam energy motif.

Experimental measurements are predicted in terms of Gibbs free energies as follows. Drug effect of 10x risdiplam ($E_{\text{ris}}$), PSI in the absence of drug ($\Psi_{\text{DMSO}}$), and PSI in the presence of 7.1x risdiplam ($\Psi_{\text{ris}}$) are predicted from $\Delta G_{\text{U1}}$ and $\Delta G_{\text{ris}}$ as in the one-interaction-mode model. Drug effect of 10x branaplam, as measured by RNA-seq followed by allelic manifold inference, is predicted using

$$E_{\text{bran}} = 1 + e^{[-\Delta G_{\text{ris}}(x)+\Delta\mu]/RT} + e^{[-\Delta G_{\text{bran}}(x)+\Delta\mu]/RT}. \tag{23}$$

PSI in the presence of 7.1x branaplam, as measured by MPSA, is predicted using

$$\Psi_{\text{bran}} = 100 \times \frac{e^{-\Delta G_{\text{U1}}(x,z)/RT} + e^{-[\Delta G_{\text{U1}}(x,z)+\Delta G_{\text{ris}}(x)]/RT} + e^{-[\Delta G_{\text{U1}}(x,z)+\Delta G_{\text{bran}}(x)]/RT}}{1 + e^{-\Delta G_{\text{U1}}(x,z)/RT} + e^{-[\Delta G_{\text{U1}}(x,z)+\Delta G_{\text{ris}}(x)]/RT} + e^{-[\Delta G_{\text{U1}}(x,z)+\Delta G_{\text{bran}}(x)]/RT}}. \tag{24}$$

We note that this model is consistent with the allelic manifold model illustrated in **Fig. 2** and **Fig. S5**. $\Delta G_{\mathrm{U1}}$ is the same in both models, and the relevant values of $\Delta G_{\mathrm{drug}}$ are given in terms of $\Delta G_{\mathrm{ris}}$ and $\Delta G_{\mathrm{bran}}$ by

$$\Delta G_{\mathrm{drug}}(x, y_{\mathrm{ris}}) = \Delta G_{\mathrm{ris}}(x), \qquad \Delta G_{\mathrm{drug}}(x, y_{\mathrm{bran}}) = \Delta G_{\mathrm{bran}}(x), \tag{25}$$

where $y_{\mathrm{ris}}$ denotes the presence of 10x risdiplam and $y_{\mathrm{bran}}$ denotes the presence of 10x branaplam.

The one-interaction-mode model for drug effect thus has 351 parameters that were inferred from MPSA and RNA-seq data: 285 parameters $\theta_{xz}^{\mathrm{U1}}$ (one context $z$, representing the *SMN2* minigene, and 285 variant 5'ss sequences $x$), 32 additive parameters $\theta_{l:b}^{\mathrm{ris}}$, 32 additive parameters $\theta_{l:b}^{\mathrm{bran}}$, 1 constant parameter $\theta_0^{\mathrm{ris}}$, and 1 constant parameter $\theta_0^{\mathrm{bran}}$. The model also has 5 hyperparameters describing experimental noise; see **Sec. 4.3** for details.

##### 3.4 Empirical concentration-dependent drug-effect model

The empirical model for concentration-dependent drug effect (**Fig. 5C,D**) is equivalent to a four-state thermodynamic model (**Fig. S11**) that generalizes the thermodynamic drug-effect model (**Fig. S5**). The pre-mRNA states and corresponding Gibbs free energies assumed by the model are as follows:

- In state 1 (Gibbs free energy 0), pre-mRNA is not bound by U1 or by drug.
- In state 2 (Gibbs free energy $\Delta G_{\mathrm{U1}}$), pre-mRNA is bound by U1 but not by drug.
- In state 3 (Gibbs free energy $H\Delta G_{\mathrm{drug}}^{0}$), pre-mRNA is bound by $H$ molecules of drug but not by U1.
- In state 4 (Gibbs free energy $\Delta G_{\mathrm{U1}} + H\Delta G_{\mathrm{drug}}^{\mathrm{U1}}$), pre-mRNA is bound by U1 and by $H$ molecules of drug.

The Gibbs free energies of the model are given in terms of the parameters $[\mathrm{drug}]$, $S$, $\mathrm{EC}_{2\mathrm{x}}$, $E_{\mathrm{max}}$, and $H$ as

$$\Delta G_{\mathrm{U1}} = -RT\log S, \qquad \Delta G_{\mathrm{drug}}^{\mathrm{U1}} = -RT\log\frac{[\mathrm{drug}]}{\mathrm{EC}_{2\mathrm{x}}}, \qquad \Delta G_{\mathrm{drug}}^{0} = -RT\log\frac{[\mathrm{drug}]}{\mathrm{EC}_{2\mathrm{x}}E_{\mathrm{max}}^{1/H}}. \tag{26}$$

Consequently, the quantities $S$, $E$, and $\Psi$ are given by:

$$\Psi = 100\times\frac{e^{-\Delta G_{\mathrm{U1}}/RT} + e^{-[\Delta G_{\mathrm{U1}}+H\Delta G_{\mathrm{drug}}^{\mathrm{U1}}]/RT}}{1+e^{-H\Delta G_{\mathrm{drug}}^{0}/RT} + e^{-\Delta G_{\mathrm{U1}}/RT} + e^{-[\Delta G_{\mathrm{U1}}+H\Delta G_{\mathrm{drug}}^{\mathrm{U1}}]/RT}} = 100\times\frac{SE}{1+SE}, \tag{27}$$

where

$$S = e^{-\Delta G_{\mathrm{U1}}/RT}, \qquad \mathrm{E} = \frac{1+e^{-H\Delta G_{\mathrm{drug}}^{\mathrm{U1}}/RT}}{1+e^{-H\Delta G_{\mathrm{drug}}^{0}/RT}} = \frac{1+\left(\frac{[\mathrm{drug}]}{\mathrm{EC}_{2\mathrm{x}}}\right)^{H}}{1+\frac{1}{E_{\mathrm{max}}}\left(\frac{[\mathrm{drug}]}{\mathrm{EC}_{2\mathrm{x}}}\right)^{H}}. \tag{28}$$

However, this thermodynamic model makes two non-physical assumptions. First is the assumption that, if any drug binds, then exactly $H$ molecules of drug bind, where $H$ is not necessarily an integer. Indeed, when fitting the empirical model to dose-response data we often observe non-integer values for $H$ (**Figs. 5E-L**, **Figs. 6A-D**). Second is the assumption that drug binds pre-mRNA in the absence of U1. This state seems implausible under the bulge-repair mechanism, but is nevertheless needed for the predicted dose-response curves to saturate at high drug concentrations. Indeed, the drug effect at saturation is given in terms of $H\Delta G_{\mathrm{drug}}^{0}$, the energy of the non-physical state, by

$$E_{\mathrm{max}} = e^{H(\Delta G_{\mathrm{drug}}^{0}-\Delta G_{\mathrm{drug}}^{\mathrm{U1}})/RT}. \tag{29}$$

We believe the non-physical aspects of this thermodynamic model, i.e., anomalous cooperativity ($H > 1$) and saturation ($\Delta G_{\mathrm{drug}}^{0} < \infty$), likely reflect biophysical mechanisms that are more complex than a simple thermodynamic model can capture. Anomalous cooperativity may arise from kinetic proofreading mechanisms that effectively read out the presence of drug bound to the U1/5'ss complex multiple times. Saturation may arise from inefficiencies in the spliceosome cycle that occur downstream of initial U1 binding to pre-mRNA (i.e., downstream of E complex formation). Nevertheless, the empirical model for concentration-dependent drug effect is useful, as it allows one to separate the drug-independent effects of pre-mRNA sequence on splicing (as quantified by $S$) from the drug-dependent effects of pre-mRNA sequence on splicing (as quantified by $H$, $\mathrm{EC}_{2\mathrm{x}}$, and $E_{\mathrm{max}}$).

Finally, we note that there are multiple reasons for parameterizing the empirical model for concentration-dependent drug effect using $\mathrm{EC}_{2\mathrm{x}}$ rather than the more conventional quantity $\mathrm{EC}_{50}$, which is defined to be the concentration of drug at which PSI is halfway between its basal value and saturation value. First, the Gibbs free energies of the equivalent biophysical model have simple expressions in terms of $\mathrm{EC}_{2\mathrm{x}}$; this is not the case for $\mathrm{EC}_{50}$. Second $\mathrm{EC}_{2\mathrm{x}}$ can be precisely inferred from dose-response data even in the absence of saturation; this is not the case for $\mathrm{EC}_{50}$. This second point is important because, in many cases, drug toxicity prevents saturating drug concentrations from being assayed. Third, the value of $\mathrm{EC}_{2\mathrm{x}}$ is not impacted by the value of $\Delta G_{\mathrm{U1}}$; this is not the case for $\mathrm{EC}_{50}$. This third point matters because $\Delta G_{\mathrm{U1}}$ is the sole Gibbs free energy in the equivalent biophysical model (**Fig. S11**) that determines context strength $S$. Biological factors that influence context strength $S$, and thus $\Delta G_{\mathrm{U1}}$, are therefore not expected to impact $\mathrm{EC}_{2\mathrm{x}}$, whereas they are expected to impact $\mathrm{EC}_{50}$. The intrinsic interdependence of $S$ and $\mathrm{EC}_{50}$ thus makes $\mathrm{EC}_{50}$ a problematic quantity to use when characterizing splice-modifying drugs. In particular, the main text argues that a failure to account for the influence of context strength on $\mathrm{EC}_{50}$ is what first led researchers[11,12] to misinterpret dose-response data as supporting the two-site hypothesis for risdiplam.

#### 4 *Bayesian model inference*

Bayesian models for allelic manifolds describing drug effect (**Fig. 2**) were defined and analyzed in STAN. Bayesian models for 5'ss-specific drug effect [the two-interaction-mode model (**Fig. 3**) and one-interaction-mode model (**Fig. S6**)] were defined and analyzed in numpyro. Bayesian models for concentration-dependent drug effect (**Fig. 5**) were defined in numpyro. For all Bayesian models, posterior parameter values were sampled using the No-U-Turn Sampler[13], which is a type of Hamiltonian Monte Carlo sampler. Reported parameter values reflect the medians and 95% credible intervals of these posterior samples.

##### 4.1 Biophysical allelic-manifold model

Drug effect values $E$ were inferred from RNA-seq read count values reported by rMATS using a simplified version of the Bayesian model described in ref.[14]. Let $k_{ij}$ denote the number of reads supporting inclusion of cassette exon $i$ in RNA sample $j$. Our Bayesian model assumes that $k_{ij}$ is sampled according to a binomial distribution,

$$k_{ij} \sim \mathrm{Binomial}\left(n_{ij}, \Phi_{ij}\right), \tag{30}$$

where $n_{ij}$ denotes the total number of experimentally observed isoform-dependent reads corresponding to cassette exon $i$ in RNA sample $j$, and $\Phi_{ij}$ is given by the PSI $\Psi_{ij}$ of cassette exon $i$ in RNA sample $j$ reweighted by the number of possible isoform-distinguishing reads for exon inclusion ($l_i^{\mathrm{inc}}$) and for exon skipping ($l_i^{\mathrm{skp}}$), i.e.,

$$\Phi_{\mathrm{ij}} = \frac{l_i^{\mathrm{inc}}\,\Psi_{ij}}{l_i^{\mathrm{inc}}\,\Psi_{ij} + l_i^{\mathrm{skp}}\,(1-\Psi_{ij})}. \tag{31}$$

The logit transform of PSI, denoted $Y_{ij}$ and related to $\Psi_{ij}$ via

$$\Psi_{\mathrm{ij}} = 100 \times \frac{\exp Y_{ij}}{1+\exp Y_{ij}}, \tag{32}$$

is further assumed to depend on the context strength of cassette exon $i$ (denoted $S_i$) and the drug effect observed for 5'ss $k$ in treatment group $m$ (denoted $E_{km}$) via

$$Y_{ij} = \log S_i + \sum_{k,m} \mathrm{X}_{jm} Z_{ik} \log E_{km} + \epsilon_{ij}\,, \tag{33}$$

where $X_{jm}$ is 1 if RNA sample $j$ corresponds to treatment condition $m$ and is zero otherwise, $Z_{ik}$ is 1 if cassette exon $i$ has 5'ss sequence $k$ and is zero otherwise, and $\epsilon_{ij}$ represents biological noise. Exon-specific context strength, $S_i$, was assigned the prior

$$\log S_i \sim \mathrm{Normal}(\alpha_k, \tau_k^2), \tag{34}$$

where $k$ is the 5'ss sequence of exon $i$, and the hyperparameters $\alpha_k$ and $\tau_k$ have the priors

$$\alpha_k \sim \mathrm{Normal}(0, 2^2), \tag{35}$$

$$\tau_k \sim \mathrm{Gamma}(2{,}0.5). \tag{36}$$

Treatment-specific and 5'ss-sequence-specific drug effect, $E_{km}$, was assigned the prior

$$\log E_{km} \sim \mathrm{Normal}(\alpha_k, \tau_k^2). \tag{37}$$

Biological noise $\epsilon_{ij}$ was assigned the prior

$$\epsilon_{ij} \sim \mathrm{Normal}(0, \sigma_k^2), \tag{38}$$

$$\sigma_k \sim \mathrm{Gamma}(2{,}0.5). \tag{39}$$

The model was coded in STAN. Posterior parameter values were sampled using the No-U-Turn Sampler[13]. Due to the model's separable structure, inference was performed separately for each 5'ss sequence $k$.

##### 4.2 Biophysical two-interaction-mode model

Let $i$ index variant 5'ss sequences; let $\Psi_i^{\text{ris,MPSA}}$, $\Psi_i^{\text{bran,MPSA}}$, and $\Psi_i^{\text{DMSO,MPSA}}$ denote PSI values measured by MPSA in the presence of 7.1x risdiplam, 7.1x branaplam, and DMSO; let $\text{E}_i^{\text{ris,RNAseq}}$ and $\text{E}_i^{\text{bran,RNAseq}}$ denote drug effect values, inferred from RNA-seq data, for 10x risdiplam and 10x branaplam; let $\theta_{lc}^{\text{ris}}$ and $\theta_{lc}^{\text{hyp}}$ denote the contribution to the risdiplam energy motif score and the hyper-activation energy motif score from base $c$ (A,C,G, or U) at nucleotide position $l$ (-4 to +6); and let $\theta_0^{\text{ris}}$ and $\theta_0^{\text{hyp}}$ denote the baseline risdiplam energy motif score and the hyper-activation energy motif score. We used Bayesian inference to estimate the parameters of the risdiplam energy motif ($\theta_{lc}^{\text{ris}}$ and $\theta_0^{\text{ris}}$) and hyper-activation energy motif ($\theta_{lc}^{\text{hyp}}$ and $\theta_0^{\text{hyp}}$) based on experimentally measured values $\Psi_i^{\text{ris,MPSA}}$, $\Psi_i^{\text{bran,MPSA}}$, $\Psi_i^{\text{DMSO,MPSA}}$, $\text{E}_i^{\text{ris,RNAseq}}$, and $\text{E}_i^{\text{bran,RNAseq}}$. The Bayesian model was defined by the equations,

$$\log \Psi_i^{\text{ris,MPSA}} = \log \Psi_i^{\text{ris,7.1x}} + \epsilon_i^{\text{ris,MPSA}}, \tag{40}$$

$$\log \Psi_i^{\text{bran,MPSA}} = \log \Psi_i^{\text{bran,7.1x}} + \epsilon_i^{\text{bran,MPSA}}, \tag{41}$$

$$\log \Psi_i^{\text{DMSO,MPSA}} = \log \Psi_i^{\text{DMSO}} + \epsilon_i^{\text{DMSO,MPSA}}, \tag{42}$$

$$\log \text{E}_i^{\text{ris,RNAseq}} = \log \text{E}_i^{\text{ris,10x}} + \epsilon_i^{\text{ris,RNAseq}}, \tag{43}$$

$$\log \text{E}_i^{\text{bran,RNAseq}} = \log \text{E}_i^{\text{bran,10x}} + \epsilon_i^{\text{bran,RNAseq}}, \tag{44}$$

$$\Psi_i^{\text{ris,7.1x}} = 100 \times \frac{S_i E_i^{\text{ris,7.1x}}}{1 + S_i E_i^{\text{ris,7.1x}}}, \tag{45}$$

$$\Psi_i^{\text{bran,7.1x}} = 100 \times \frac{S_i E_i^{\text{bran,7.1x}}}{1 + S_i E_i^{\text{bran,7.1x}}}, \tag{46}$$

$$\Psi_i^{\text{DMSO}} = 100 \times \frac{S_i}{1 + S_i}, \tag{47}$$

$$E_{\text{i}}^{\text{ris,7.1x}} = 1 + \exp[\phi_i^{\text{ris}}], \tag{48}$$

$$E_{\text{i}}^{\text{ris,10x}} = 1 + \exp[\phi_i^{\text{ris}} + \Delta\mu], \tag{49}$$

$$E_{\text{i}}^{\text{bran,7.1x}} = 1 + \exp[\phi_i^{\text{ris}}] + \exp[\phi_i^{\text{hyp}}] \tag{50}$$

$$E_{\text{i}}^{\text{bran,10x}} = 1 + \exp[\phi_i^{\text{ris}} + \Delta\mu] + \exp[\phi_i^{\text{hyp}} + \Delta\mu] \tag{51}$$

$$\phi_i^{\text{ris}} = \theta_0^{\text{ris}} + \sum_{l,c} \theta_{lc}^{\text{ris}} x_{ilc}, \tag{52}$$

$$\phi_i^{\text{hyp}} = \theta_0^{\text{hyp}} + \sum_{l,c} \theta_{lc}^{\text{hyp}} x_{ilc}, \tag{53}$$

and by the priors,

$$\epsilon_i^{\text{ris,MPSA}} \sim \text{Normal}\left(0, \sigma_{\text{MPSA}}^{\text{ris}}\right), \tag{54}$$

$$\epsilon_i^{\text{bran,MPSA}} \sim \text{Normal}\left(0, \sigma_{\text{MPSA}}^{\text{bran}}\right), \tag{55}$$

$$\epsilon_i^{\text{DMSO,MPSA}} \sim \text{Normal}\left(0, \sigma_{\text{MPSA}}^{\text{DMSO}}\right), \tag{56}$$

$$\epsilon_i^{\text{ris,RNAseq}} \sim \text{Normal}\left(0, \sigma_{\text{RNAseq}}^{\text{ris}}\right), \tag{57}$$

$$\epsilon_i^{\text{bran,MPSA}} \sim \text{Normal}\left(0, \sigma_{\text{RNAseq}}^{\text{bran}}\right), \tag{58}$$

$$\sigma_{\text{MPSA}}^{\text{ris}}, \sigma_{\text{MPSA}}^{\text{bran}}, \sigma_{\text{MPSA}}^{\text{DMSO}}, \sigma_{\text{RNAseq}}^{\text{ris}}, \sigma_{\text{RNAseq}}^{\text{bran}} \sim \text{Exponential}(1), \tag{59}$$

$$\theta_0^{\mathrm{ris}}, \theta_{lc}^{\mathrm{ris}}, \theta_0^{\mathrm{hyp}}, \theta_{lc}^{\mathrm{hyp}} \sim \mathrm{Normal}(0, 5^2), \tag{60}$$

$$\log S_j \sim \mathrm{Normal}(0,5^2). \tag{61}$$

The model was coded in numpyro. Posterior parameter values were sampled using the No-U-Turn Sampler[13]. The sequence logo and scatter plot in **Fig. 3G** show mean-centered values of $\theta_{lc}^{\mathrm{ris}}$, i.e., $\Delta\Delta G_{lc}^{\mathrm{ris}} = \theta_{lc}^{\mathrm{ris}} - \frac{1}{4}\sum_{c'} \theta_{lc'}^{\mathrm{ris}}$. Similarly, **Fig. 3H** shows mean-centered values of $\theta_{lc}^{\mathrm{hyp}}$, i.e., $\Delta\Delta G_{lc}^{\mathrm{hyp}} = \theta_{lc}^{\mathrm{hyp}} - \frac{1}{4}\sum_{c'} \theta_{lc'}^{\mathrm{hyp}}$. This mean-centering procedure is necessary to remove non-identifiable degrees of freedoms (called "gauge freedoms") from each posterior sample.

##### 4.3 Biophysical one-interaction-mode model

The one-interaction-mode model was inferred in the same manner as the two-interaction-mode model (**Sec. 4.2**), but replacing **Eq. 50, Eq. 51,** and **Eq. 53** with

$$E_{\mathrm{i}}^{\mathrm{bran,7.1x}} = 1 + \exp[\phi_i^{\mathrm{bran}}], \tag{62}$$

$$E_{\mathrm{i}}^{\mathrm{bran,10x}} = 1 + \exp[\phi_i^{\mathrm{bran}} + \Delta\mu], \tag{63}$$

$$\phi_i^{\mathrm{bran}} = \theta_0^{\mathrm{bran}} + \sum_{l,c} \theta_{lc}^{\mathrm{bran}} x_{ilc}, \tag{64}$$

And replacing $\theta_{lc}^{\mathrm{hyp}}$ and $\theta_0^{\mathrm{hyp}}$ in **Eq. 59** with $\theta_{lc}^{\mathrm{bran}}$ and $\theta_0^{\mathrm{bran}}$, the parameters of the branaplam energy motif. The sequence logo and scatter plot in **Fig. S6F** show mean-centered values of $\theta_{lc}^{\mathrm{ris}}$, i.e., $\Delta\Delta G_{lc}^{\mathrm{ris}} = \theta_{lc}^{\mathrm{ris}} - \frac{1}{4}\sum_{c'} \theta_{lc'}^{\mathrm{ris}}$. Similarly, **Fig. S6G** shows mean-centered values of $\theta_{lc}^{\mathrm{bran}}$, i.e., $\Delta\Delta G_{lc}^{\mathrm{bran}} = \theta_{lc}^{\mathrm{bran}} - \frac{1}{4}\sum_{c'} \theta_{lc'}^{\mathrm{bran}}$. Note that this one-interaction-mode model has the same number of parameters as the two-interaction-mode model.

##### 4.4 Empirical concentration-dependent drug-effect model

Empirical models of concentration-dependent drug effect were fit to dose-response data as follows. Let $i$ index samples, including biological replicates and samples treated with different drug concentrations; let $C_i$ denote the concentration of drug used for sample $i$ by RT-qPCR; let $R_i^{\mathrm{qPCR}}$ denote the inclusion/exclusion ratio measured for sample $i$; let $R_i$ denote the model-predicted inclusion/exclusion ratio; and let $E_i$ denote the model-predicted drug effect at drug concentration $C_i$. The Bayesian empirical model of concentration-dependent drug effect was defined by,

$$\log R_{\mathrm{i}}^{\mathrm{qPCR}} = \log R_{\mathrm{i}} + \epsilon_{\mathrm{i}}, \tag{65}$$

$$R_i = S E_i, \tag{66}$$

$$E_i = \frac{1 + \left(\frac{C_i}{\mathrm{EC}_{2\mathrm{x}}}\right)^H}{1 + \frac{1}{E_{\max}}\left(\frac{C_i}{\mathrm{EC}_{2\mathrm{x}}}\right)^H}, \tag{67}$$

and by the priors,

$$\log_{10} S \sim \mathrm{Uniform}(-3, 3), \tag{68}$$

$$\epsilon_{\mathrm{i}} \sim \mathrm{Normal}(0, \sigma^2), \tag{69}$$

$$\log_2 H \sim \mathrm{Uniform}(-2, 2), \tag{70}$$

$$\log_{10} \mathrm{EC}_{2\mathrm{x}} \sim \mathrm{Uniform}(-3,3), \tag{71}$$

$$\log_{10} E_{\max} \sim \mathrm{Uniform}(0, 6), \tag{72}$$

$$\log_{10} \sigma \sim \mathrm{Uniform}(-2,2). \tag{73}$$

The model was coded in numpyro. Posterior parameter values were sampled using the No-U-Turn Sampler[13]. Each dose-response curve was analyzed separately.

##### 4.5 Quadratic linear-mixture drug-effect model

Quadratic curves were fit to linear-mixture data as follows. Let $i$ index biological samples, including biological replicates and samples treated with different mixtures of two given drugs; let $x_i$ denote the normalized concentration for drug 1, with values ranging from 0 to 1; let $R_i^{\mathrm{qPCR}}$ denote the inclusion/exclusion ratio measured for sample $i$; and let $R_i$ denote the inclusion/exclusion ratio predicted by the model at normalized drug concentration $x_i$. The Bayesian quadratic model was defined by,

$$\log_2 R_{\mathrm{i}}^{\mathrm{qPCR}} = \log_2 R_{\mathrm{i}} + \epsilon_{\mathrm{i}}, \quad (74)$$

$$\log_2 R_i \ = a + b(x - 0.5) + c(x - 0.5)^2, \quad (75)$$

and by the prior,

$$\epsilon_{\mathrm{i}} \sim \mathrm{Normal}(0, \sigma^2), \quad (76)$$

$$\log_{10} \sigma \ \sim \mathrm{Uniform}(-3,1), \quad (77)$$

$$a, b, c \sim \mathrm{Normal}(0,10^2). \quad (78)$$

The model was coded in numpyro. Posterior parameter values were sampled using the No-U-Turn Sampler[13]. Each linear-mixture curve was analyzed separately. Synergy was assessed for using p-values representing the probability (fraction of posterior samples) for which the extremum value of the quadratic model was smaller than both of the boundary values of the quadratic model.

### 5 Supplemental Tables

#### 5.1 *Table S1. Inferred parameters for dose-response curves*

Listed are inferred parameter values for the dose-response curves in **Fig. 5**, **Fig. 6**, and **Fig. S13**. Models were defined as described in **Sec. 3.1** and **Sec. 4.1**. Values indicate medians; brackets indicate 95% credible intervals. Bayesian posterior distributions were sampled using Hamiltonian Monte Carlo. $P$, p-values from **Fig. S13** for the no-synergy null hypothesis (i.e., that $H_{\mathrm{mix}}$ is not larger than both $H_{\mathrm{drug\ 1}}$ and $H_{\mathrm{drug\ 2}}$) computed by comparing posterior-sampled values of $H$ for the drug mixture to posterior-sampled values of $H$ for each individual drug. n.s., $P \geq 0.5$; ***, $P < 10^{-3}$.

| FIG. | TREATMENT | MINIGENE | $\log_{10} S$ | $log_{10} E_{max}$ | $\mathrm{EC}_{2x}$ | $H$ | VS. | $P$ |
|---|---|---|---|---|---|---|---|---|
| **5E** | risdiplam | *SMN2* WT | 0.2 [0.1, 0.2] | 4.0 [2.7, 5.7] | 16.7 [14.3, 19.2] (nM) | 1.25 [1.19, 1.32] | N/A | N/A |
| **5F** | risdiplam | *SMN2* 25T26T | -2.2 [-2.2, -2.1] | 4.9 [3.9, 6.0] | 10.2 [7.2, 13.4] (nM) | 1.61 [1.49, 1.75] | N/A | N/A |
| **5G** | risdiplam | *SMN2* Δ22-27 | -2.3 [-2.4, -2.3] | 5.0 [4.0, 6.0] | 10.9 [8.1, 14.0] (nM) | 1.67 [1.55, 1.80] | N/A | N/A |
| **5H** | risdiplam | *SMN2* Δ17-28 | -1.4 [-1.4, -1.3] | 5.0 [4.0, 6.0] | 11.0 [8.0, 14.1] (nM) | 1.74 [1.60, 1.87] | N/A | N/A |
| **5I** | branaplam | *SMN2* WT | 0.2 [0.1, 0.2] | 2.5 [2.3, 2.8] | 12.2 [10.0, 14.4] (nM) | 1.37 [1.26, 1.48] | N/A | N/A |
| **5J** | branaplam | *SMN2* 25T26T | -2.3 [-2.4, -2.3] | 4.5 [3.2, 6.0] | 16.2 [11.8, 20.8] (nM) | 1.86 [1.62, 2.09] | N/A | N/A |
| **5K** | branaplam | *SMN2* Δ22-27 | -2.3 [-2.4, -2.2] | 5.0 [3.9, 6.0] | 8.2 [6.1, 10.4] (nM) | 1.62 [1.51, 1.74] | N/A | N/A |
| **5L** | branaplam | *SMN2* Δ17-28 | -1.3 [-1.4, -1.3] | 4.6 [3.6, 6.0] | 6.2 [4.7, 8.0] (nM) | 1.46 [1.36, 1.56] | N/A | N/A |
| **6A** | ASOi7 | *SMN2* WT | 0.1 [0.1, 0.2] | 2.4 [2.3, 2.4] | 0.2 [0.1, 0.2] (nM) | 1.34 [1.26, 1.43] | N/A | N/A |
| **6B** | ASOi6 | *SMN2* WT | 0.1 [0.1, 0.1] | 1.7 [1.5, 2.0] | 0.7 [0.6, 0.9] (nM) | 1.07 [1.00, 1.17] | N/A | N/A |
| **6C** | RECTAS | *ELP1* WT | -0.2 [-0.3, -0.2] | 2.0 [1.9, 2.1] | 272.7 [226.1, 320.9] (nM) | 1.13 [1.06, 1.22] | N/A | N/A |
| **6D** | ASOi20 | *ELP1* WT | -0.2 [-0.3, -0.2] | 2.8 [2.0, 5.2] | 100.8 [71.3, 134.3] (nM) | 1.09 [1.00, 1.23] | N/A | N/A |
| **S13A** | risdiplam + branaplam | *SMN2* WT | 0.2 [0.1, 0.2] | 2.9 [2.4, 4.8] | 1.5 [1.1, 1.8] (1x) | 1.36 [1.24, 1.49] | **5E, 5I** | $5.3 \times 10^{-1}$ (n.s.) |
| **S13B** | risdiplam + ASOi6 | *SMN2* WT | 0.1 [0.1, 0.2] | 2.5 [2.4, 2.6] | 1.9 [1.5, 2.2] (1x) | 1.51 [1.41, 1.61] | **5E, 6B** | $8.8 \times 10^{-4}$ (***) |
| **S13C** | risdiplam + ASOi7 | *SMN2* WT | 0.2 [0.1, 0.2] | 2.5 [2.4, 2.6] | 1.6 [1.4, 1.9] (1x) | 1.59 [1.48, 1.70] | **5E, 6A** | $6.2 \times 10^{-4}$ (***) |
| **S13D** | branaplam + ASOi6 | *SMN2* WT | 0.1 [0.1, 0.2] | 2.3 [2.2, 2.4] | 1.8 [1.4, 2.3] (1x) | 1.46 [1.34, 1.59] | **5I, 6B** | $1.1 \times 10^{-1}$ (n.s.) |
| **S13E** | branaplam + ASOi7 | *SMN2* WT | 0.2 [0.1, 0.2] | 2.6 [2.5, 2.7] | 2.2 [1.9, 2.6] (1x) | 1.73 [1.62, 1.85] | **5I, 6A** | $1.9 \times 10^{-4}$ (***) |
| **S13F** | ASOi6 + ASOi7 | *SMN2* WT | 0.0 [-0.1, 0.1] | 2.6 [2.2, 3.5] | 3.9 [3.0, 4.8] (1x) | 1.35 [1.20, 1.47] | **6A, 6B** | $4.8 \times 10^{-1}$ (n.s.) |
| **S13G** | RECTAS + ASOi20 | *ELP1* WT | -0.0 [-0.1, 0.0] | 4.2 [2.6, 6.0] | 1.7 [1.3, 2.1] (1x) | 1.67 [1.49, 1.85] | **6C, 6D** | $1.1 \times 10^{-4}$ (***) |

#### 5.2 *Table S2. Minigene plasmids and minigene plasmid libraries*

Listed are the minigene plasmids and minigene plasmid libraries used in this study. See **Sec. 1.3** and **Sec. 1.4** for descriptions of how these plasmids and plasmid libraries were constructed.

| NAME | DESCRIPTION |
|---|---|
| **pSMN2_WT** | Primary *SMN2* minigene plasmid. Used in dose-response experiments and linear-mixture experiments (**Fig. 5E,I**, **Fig. 6A,B,E-J**, **Fig. S12E-J**, **Fig. S12E,F**, and **Fig. S13A-F**). SnapGene file is provided on GitHub. |
| **pSMN2_PT_mut1** | pSMN2_WT with PT mutation ex7:G25T,G26T. Used in **Fig. 5F,J**. |
| **pSMN2_PT_mut2** | pSMN2_WT with PT mutation ex7:Δ22..27. Used in **Fig. 5G,K**. |
| **pSMN2_PT_mut3** | pSMN2_WT with PT mutation ex7:Δ17..28. Used in **Fig. 5H,L**. |
| **pSMN2_cDNA_full** | pSMN2_WT with intron 6 and intron 7 excised. Used to validate qPCR primers for *SMN2* minigenes. |
| **pSMN2_cDNA_Δ7** | pSMN2_WT with intron 6, exon 7, and intron 7 excised. Used to validate qPCR primers for *SMN2* minigenes. |
| **pSMN2_5ss_libX (X=1,2,3)** | Minigene library created from pSMN2_WT by replacing the 5'ss of exon 7 with 285 variant 5'ss sequences (1 WT + 24 single mutants + 252 double mutants + 4 consensus + 4 null mutants). Used in *SMN2* MPSA experiments (**Fig. 1, Fig. 3, Fig. S1, Fig. S3, Fig. S4, Fig. S6, Fig. S7, Fig. S8, Fig. S9**). |
| **pSMN2_ssbc_libX (X=1,2,3)** | Precursor plasmid library of pSMN2_5ss_libX, i.e., library before insertion, between the AarI restriction sites, of the fragment containing intron 7. |
| **pSMN2_5ss_consN**<br>**(N=A,C,G,U)** | Four variants of pSMN2_WT, each of which contains a different consensus 5'ss (having the form NCAG/GUAAGU) at exon 7. N refers to the nucleotide at position -4. Used to validate the MPSA (**Fig. S1I**). |
| **pSMN2_5ss_nullN**<br>**(N=A,C,G,U)** | Four variants of pSMN2_WT, each of which contains a different null 5'ss (having the form NCAG/GGAAGU) at exon 7. N refers to the nucleotide at position -4. Used to validate the MPSA (**Fig. S1I**). |
| **pSMN1_5ss_mutX**<br>**(X=A3C, A3G, A3U, A4C, A4G, A4U, G5A, G5C, G5U)** | pSMN2_WT with an ex7:T6C mutation (which matches the *SMN1* exon 7 sequence and increases basal PSI) and with various mutations in the intronic region of the exon 7 5'ss. Used in qPCR assays to validate the specificities of risdiplam and branaplam for 5'ss intronic positions (**Fig. 5F,G**). |
| **pELP1_WT** | Primary ELP1 minigene plasmid. Used in dose-response experiments and linear-mixture experiments (**Fig. 6C,D,K, Fig. S12A-D, Fig. S13G**). SnapGene file is provided on GitHub. |
| **pELP1_cDNA_full** | pEPL1_WT with intron 19 and intron 20 excised. Used to validate qPCR primers for *ELP1* minigenes. |
| **pELP1_cDNA_Δ20** | pEPL1_WT with intron 19, exon 20, and intron 20 excised. Used to validate qPCR primers for *ELP1* minigenes. |
| **pELP1_5ss_libX (X=1,2)** | Minigene library from ref.[1], created from pELP1_WT by replacing the 5'ss of exon 7 with approximately 32,768 variant 5'ss of the form ANNN/GYNNNN. Used in *ELP1* MPSA experiments (**Fig. S2**). |
| **pELP1_ssbc_libX (X=1,2)** | Precursor of pELP1_5ss_libX, before inserting intron 20. |

#### 5.3 *Table S3. Primers and antisense oligonucleotides*

Listed are the key primers and antisense oligonucleotides (ASOs) used in this study.

| NAME | SEQUENCE | DESCRIPTION |
|---|---|---|
| **BCR_short** | 5'-GGCAACTAGAAGGCACA-3' | Reverse-transcription primer used to generate cDNA for all MPSA and qPCR experiments. |
| **SBCRmod** | 5'-AACTAGAAGGCACAGTCG-3' | Reverse primer for qPCR on *SMN2* & *ELP1* cDNA. |
| **SMN2_inclusion_fwd** | 5'-GGCTATCATACTGGCTATTATATGGGTT-3' | Forward primer for qPCR on *SMN1/2* +e7 cDNA. |
| **SMN2_exclusion_fwd** | 5'-GGCTATCATACTGGCTATTATATGGAAA-3' | Forward primer for qPCR for *SMN1/2* -e7 cDNA. |
| **ELP1_inclusion_fwd** | 5'-TCGGAAGTGGTTGGACAAACTT-3' | Forward primer for qPCR for *ELP1* +e20 cDNA. |
| **ELP1_exclusion_fwd** | 5'-GACACAAAGCTTGTATTACAGACTT-3' | Forward primer for qPCR for *ELP1* -e20 cDNA. |
| **ASO_SMN2_in7:9-23** | 5'-TTTCATAATGCTGGC-3' | ASO with PO 2'MOE chemistry targeting *SMN2* intron 7. Sequence is from ref.[15] |
| **ASO_SMN2_in6:55-41** | 5'-AGATATAGATAGCTA-3' | ASO with PO 2'MOE chemistry targeting *SMN2* intron 6. Sequence is from ref.[15] |
| **ASO_ELP1_in20:7-26** | 5'-GTCGCAAACAGTACAATGGC-3' | ASO with PO 2'MOE chemistry targeting *ELP1* intron 20. Sequence is from ref.[16] |
| **SMN2 BseRI ss top** | 5'-CATCATGAGGAGACTTAAATTANNNNGTNNNNCTGCCAGCATGCAG GTGGATGCACATGATGACATAA-3' | Oligo library of 285 sequences ordered to IDT |
| **ELP1 BseRI ss top** | 5'-CATCATGAGGAGAGTGGTTGGANNNGYNNNNGCCATTGTGCAGGT GGATGCACATGATGACATAAT-3' | Oligo library of 32,768 sequences |
| **SMN2 NotI bc bot** | 5'-ATGATGGCGGCCGCNNNNNNNNNNNNNNNNNNNNTCTAGAATGCA GGTGATTATGTCATCATGTGCATC-3' | Oligo annealed with ss top including N20 barcode |
| **ELP1 XhoI bc bot** | 5'-ATGATGCTCGAGNNNNNNNNNNNNNNNNNNNNTCTAGAATGCAGG TGATTATGTCATCATGTGCATC-3' | Oligo annealed with ss top including N20 barcode |
| **SMN2 BB fwd** | 5'-AGATCTGGAATGTGAAGCGTT-3' | Forward primer for amplifying backbone for *SMN2* ssbc plasmid construction |
| **SMN2 BB rev** | 5'-GATGATGGAGGAGTGATGAGTTAATTTAAGGAATGTG AGC-3' | Reverse primer for amplifying backbone for *SMN2* ssbc plasmid construction |
| **ELP1 BB fwd** | 5'-CATTTGAATGCATGAGAAAGCTG-3' | Forward primer for amplifying backbone for *ELP1* ssbc plasmid construction |
| **ELP1 BB rev** | 5'-GATGATGGAGGAGTGATGAGTTCCAACCACTTCCGAAT CTGAG-3' | Reverse primer for amplifying backbone for *ELP1* ssbc plasmid construction |
| **SMN2 PE1 top** | 5'-AATGATACGGCGACCACCGAGATCTACACTCTTTCCCTACACGACG CTCTTCCGATCTNNNNNNNNNNGTGCT-3' | Primer to ligate against digested *SMN2* ssbc plasmid, adapter 1 top strand |
| **SMN2 PE1 bot** | 5'-CNNNNNNNNNNAGATCGGAAGAGCGTCGTGTAGGGAAAGAGTGTA GATCTCGGTGGTCGCCGTATCATT-3' | Primer to ligate against digested *SMN2* ssbc plasmid, adapter 1 bottom strand |
| **SMN2 PE2 top** | 5'-GGCCGCNNNNNNNNNNAGATCGGAAGAGCGGTTCAGCAGGAATG CCGAGACCGATCTCGTATGCCGTCTTCTGCTT-3' | Primer to ligate against digested *SMN2* ssbc plasmid, adapter 2 top strand |
| **SMN2 PE2 bot** | 5'-AAGCAGAAGACGGCATACGAGATCGGTCTCGGCATTCCTGCTGAA CCGCTCTTCCGATCTNNNNNNNNNNGC-3' | Primer to ligate against digested *SMN2* ssbc plasmid, adapter 2 bottom strand |
| **ELP1 PE1 top** | 5'-AATGATACGGCGACCACCGAGATCTACACTCTTTCCCTACACGACG CTCTTCCGATCTNNNNNNNNNNTTAAT-3' | Primer to ligate against digested *ELP1* ssbc plasmid, adapter 1 top strand |
| **ELP1 PE1 bot** | 5'-TAANNNNNNNNNNAGATCGGAAGAGCGTCGTGTAGGGAAAGAGTG TAGATCTCGGTGGTCGCCGTATCATT-3' | Primer to ligate against digested *ELP1* ssbc plasmid, adapter 1 bottom strand |
| **ELP1 PE2 top** | 5'-TCGAGNNNNNNNNNNAGATCGGAAGAGCGGTTCAGCAGGAATGCC GAGACCGATCTCGTATGCCGTCTTCTGCTT-3' | Primer to ligate against digested *ELP1* ssbc plasmid, adapter 2 top strand |
| **ELP1 PE2 bot** | 5'-AAGCAGAAGACGGCATACGAGATCGGTCTCGGCATTCCTGCTGAAC CGCTCTTCCGATCTNNNNNNNNNNC-3' | Primer to ligate against digested *ELP1* ssbc plasmid, adapter 2 bottom strand |
| **SMN2 insert F** | 5'-CATCATCACCTGCTAGGGCCAGCATTATGAAACTGAATC-3' | Forward primer for amplifying *SMN2* intron 7 to insert in *SMN2* ssbc AarI sites |
| **SMN2 insert R** | 5'-ATGATGCACCTGCCCTATCTAGAATAACGCTTCACATTCCAGATC-3' | Reverse primer for amplifying *SMN2* intron 7 to insert in *SMN2* ssbc AarI sites |
| **ELP1 insert F** | 5'-CATCGTCACCTGCAAGCGCCATTGTACTGTTTGCGACTAGTTAGC-3' | Forward primer for amplifying *ELP1* intron 20 to insert in *ELP1* ssbc AarI sites |
| **ELP1 insert R** | 5'-ATGATGCACCTGCCATGTCTAGAACTACTTAGGGTTATGATCAT-3' | Reverse primer for amplifying ELP1 intron 20 to insert in *ELP1* ssbc AarI sites |

| | | |
|---|---|---|
| **SMN2 e7F** | 5'-GAAGGAAGGTGCTCACATTC-3' | Forward primer for *SMN2* MPSA inclusion isoform amplification |
| **ELP1 e20F** | 5'-GTTGTTCATCATCGAGCCCTGG-3' | Forward primer for *ELP1* MPSA inclusion isoform amplification |
| **SMN2 e8F** | 5'-GACACCACTAAAGAAACGATCAG-3' | Forward primer for *SMN2* MPSA barcode amplification |
| **ELP1 e21F** | 5'-GCATGAGAAAGCTGAGAATC-3' | Forward primer for *ELP1* MPSA barcode amplification |
| **BCR** | 5'-GGCAACTAGAAGGCACAGTCG-3' | Reverse primer for inclusion isoform and barcode amplification |
| **SMN2 barcode-LID F** | 5'-CCCTACACGACGCTCTTCCGATCTNNNNNNNNNNGACACCACTAAAGAAACGATCAG-3' | Forward primer for *SMN2* sample-specific barcode addition |
| **SMN2 barcode-LID R** | 5'-CATTCCTGCTGAACCGCTCTTCCGATCTNNNNNNNNNNGGCAACTAGAAGGCACAGTCG-3' | Reverse primer for *SMN2* sample-specific barcode addition |
| **ELP1 barcode-LID F** | 5'-CCCTACACGACGCTCTTCCGATCTNNNNNNNNNGCATGAGAAAGCTGAGAATC-3' | Forward primer for *ELP1* sample-specific barcode addition |
| **ELP1 barcode-LID R** | 5'-CATTCCTGCTGAACCGCTCTTCCGATCTNNNNNNNNNNGGCAACTAGAAGGCACAGTCG-3' | Reverse primer for *ELP1* sample-specific barcode addition |
| **PE1_v4** | 5'-AATGATACGGCGACCACCGAGATCTACACTCTTTCCCTACACGACGCTCTTC-3' | Forward primer for Illumina-compatible end addition |
| **PE2_v4** | 5'-AAGCAGAAGACGGCATACGAGATCGGTCTCGGCATTCCTGCTGAACCGCT-3' | Reverse primer for Illumina-compatible end addition |
| **SMN2 e6F** | 5'-GGGAAGTATGTTAATTTCATGGTACATGAGTGG-3' | Forward primer for Radioactive gel assay |

#### *5.4 Table S4. Key resources*

Listed are the key reagents used in this study.

| REAGENTS OR RESOURCE | SOURCE | IDENTIFIER |
|---|---|---|
| **Risdiplam** | MedChemExpress | HY-109101 (lot # 114275; 98.15% pure) |
| **Branaplam** | MedChemExpress | HY-19620 (lot # 79979; 98.04% pure) |
| **RECTAS** | J&W Pharmlab, LLC | 70R0445 (lot # JWY551-220, 98% pure) |
| **HeLa cells** | CSHL cell line repository | N/A |
| **15-cm plates** | Corning | 430599 |
| **12-well plates** | Falcon | 353043 |
| **Lipofectamine™ 2000 Transfection Reagent** | ThermoFisher Scientific | 11668-019 |
| **Opti-MEM™ Reduced Serum Medium** | ThermoFisher Scientific | 31985088 |
| **Dulbeco's Modification of Eagle's Medium (DMEM)** | Corning | 10-013-CV |
| **DMSO** | Millipore Sigma | D2650-5X5ML |
| **TRIzol™ Reagent** | ThermoFisher Scientific | 15596018 |
| **PowerUP™ SYBR™ Green Master Mix** | ThermoFisher Scientific | A25742 |
| **ROX reference dye** | ThermoFisher Scientific | 12223012 |
| **Antisense oligonucleotides** | Integrated DNA Technologies (IDT) | N/A |
| **Primers** | Integrated DNA Technologies (IDT) and Sigma | N/A |
| **Illumina high-throughput sequencing** | CSHL NextGen DNA sequencing core facility | N/A |

#### 6 Supplemental Figures

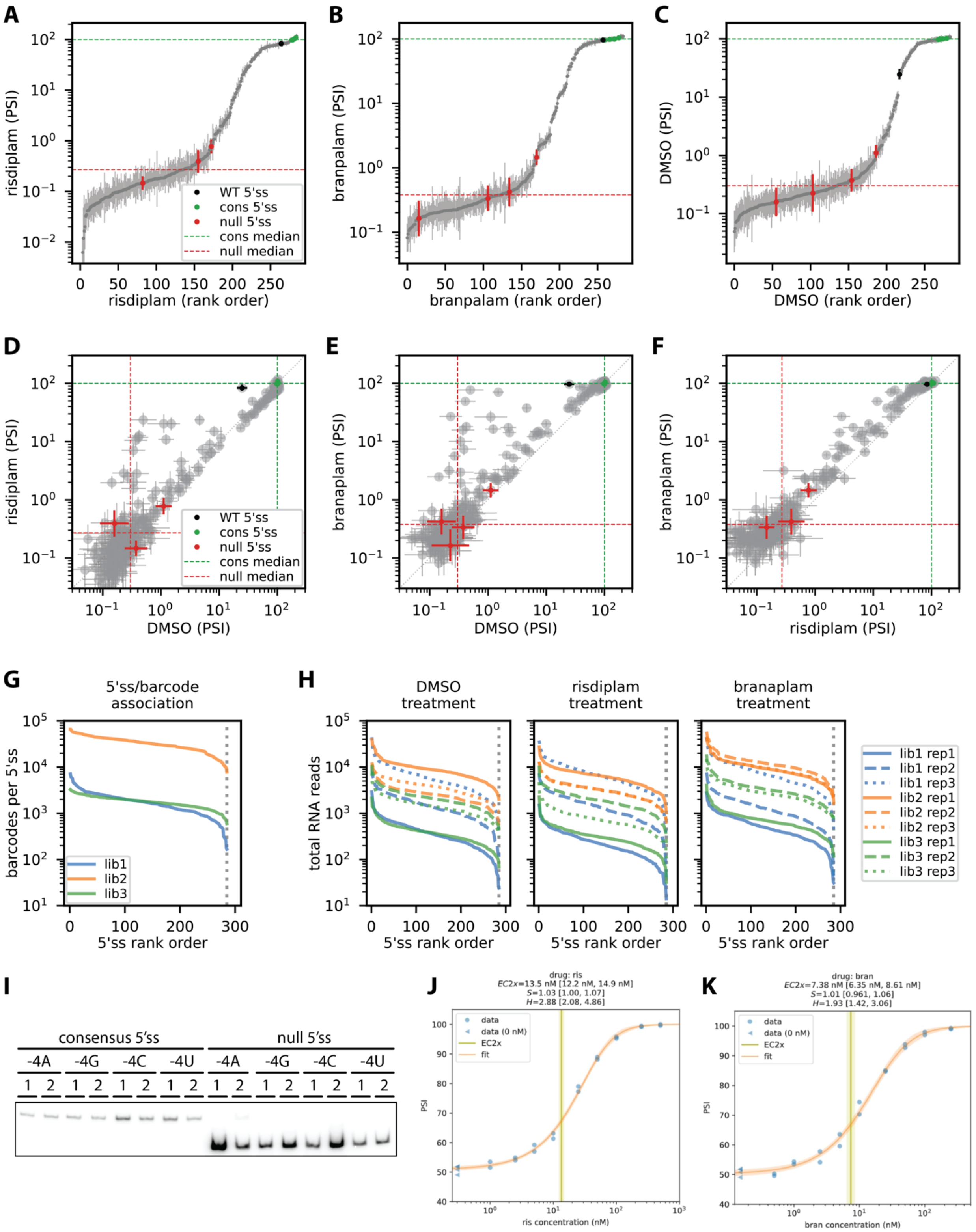

##### *6.1 Figure S1. Dynamic range and precision of MPSA data*

MPSA data for the *SMN2* exon 7 minigene library. **(A-C)** Dynamic range and precision of PSI values for cells treated with (**A**) risdiplam, (**B**) branaplam, or (**C**) DMSO. **(D-F)** Same data as in **A-C**, but showing (**D**) risdiplam versus DMSO, (**E**) branaplam versus DMSO, or (**F**) branaplam versus risdiplam. Dots, median PSI. Error bars, 95% confidence interval on PSI. Black dots, wild-type *SMN2* exon 7 5'ss (AGGA/GUAAGU). Green dots, consensus 5'ss sequences (NCAG/GUAAGU). Red dots, null 5'ss sequences (NCAG/GGAAGA; the G at position +2 abrogates usage as a 5’ss). Green dashed line, median value of consensus 5’ss over all biological replicates. Red dashed line, median value of null 5’ss over all biological replicates. (**G**) Number of 5’ss associated with each barcode in each (independently cloned) *SMN2* sub-library (lib1, lib2, lib3). (**H**) Number of total isoform reads obtained for each 5’ss in each replicate (rep1, rep2, rep3) of each *SMN2* sub-library in each treatment condition (DMSO, risdiplam, branaplam). Vertical dotted line indicates the expected number of variant 5’ss (i.e., 285). (**I**) Radioactive RT-PCR gels for the four consensus 5’ss and four null 5’ss (two biological replicates per 5’ss). MPSA, massively parallel splicing assay. 5’ss, 5’ splice site. (**J,K**) Pilot radioactive RT-PCR dose-response experiments for (**J**) risdiplam and (**K**) branaplam in the *SMN2* minigene context. MPSA, massively parallel splicing assay; PSI, percent spliced in; DMSO, dimethyl sulfoxide; 5’ss, 5’ splice site.

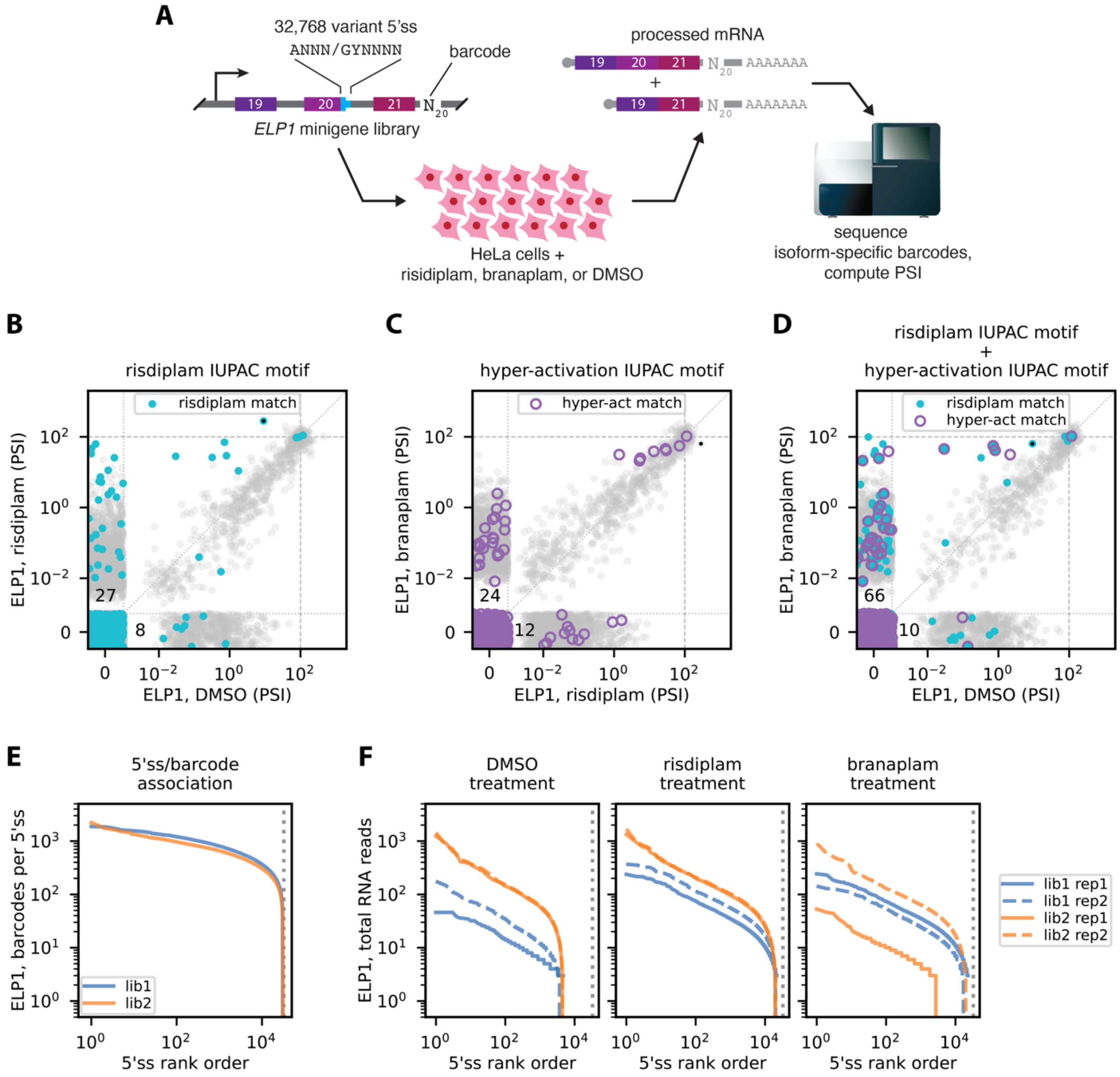

##### *6.2 Figure S2. IUPAC motif performance on MPSA data for the* **ELP1** *minigene library*

(**A**) MPSA performed in the context of a minigene library (reported previously[1]) containing exons 19, 20, and 21 of *ELP1*. In this library, the 5'ss of exon 20 was replaced by approximately 32,768 variant 5'ss sequences of the form ANNN/GYNNNN. (**B-D**) PSI values measured in the presence of (**B**) risdiplam vs. DMSO, (**C**) branaplam vs. risdiplam, or (**D**) branaplam vs. DMSO. The predictions of the risdiplam IUPAC motif (from **Fig. 1D**) and the hyper-activation IUPAC motif (from **Fig. 1E**) are also shown. PSI values of 100 are indicated by dashed lines. PSI values to the left of the vertical dotted line or below the horizontal dotted line are equal to zero; random jitter has been added to aid visualization. Numbers quantify motif-matched points within corresponding sections of plot (x=0 and y>0, or x>0 and y=0). Black dot, wild-type *SMN2* exon 7 5'ss (AGGA/GUAAGU). (**E**) Number of 5'ss associated with each barcode in each (independently cloned) *ELP1* sub-library (lib1, lib2). (**F**) Number of total isoform reads obtained for each 5'ss in each replicate (rep1, rep2) of each *ELP1* sub-library in each treatment condition (DMSO, risdiplam, branaplam). Vertical dotted line indicates the expected number of variant 5'ss (i.e., 32,768). MPSA, massively parallel splicing assay; DMSO, dimethyl sulfoxide; 5'ss, 5' splice site; PSI, percent spliced in.

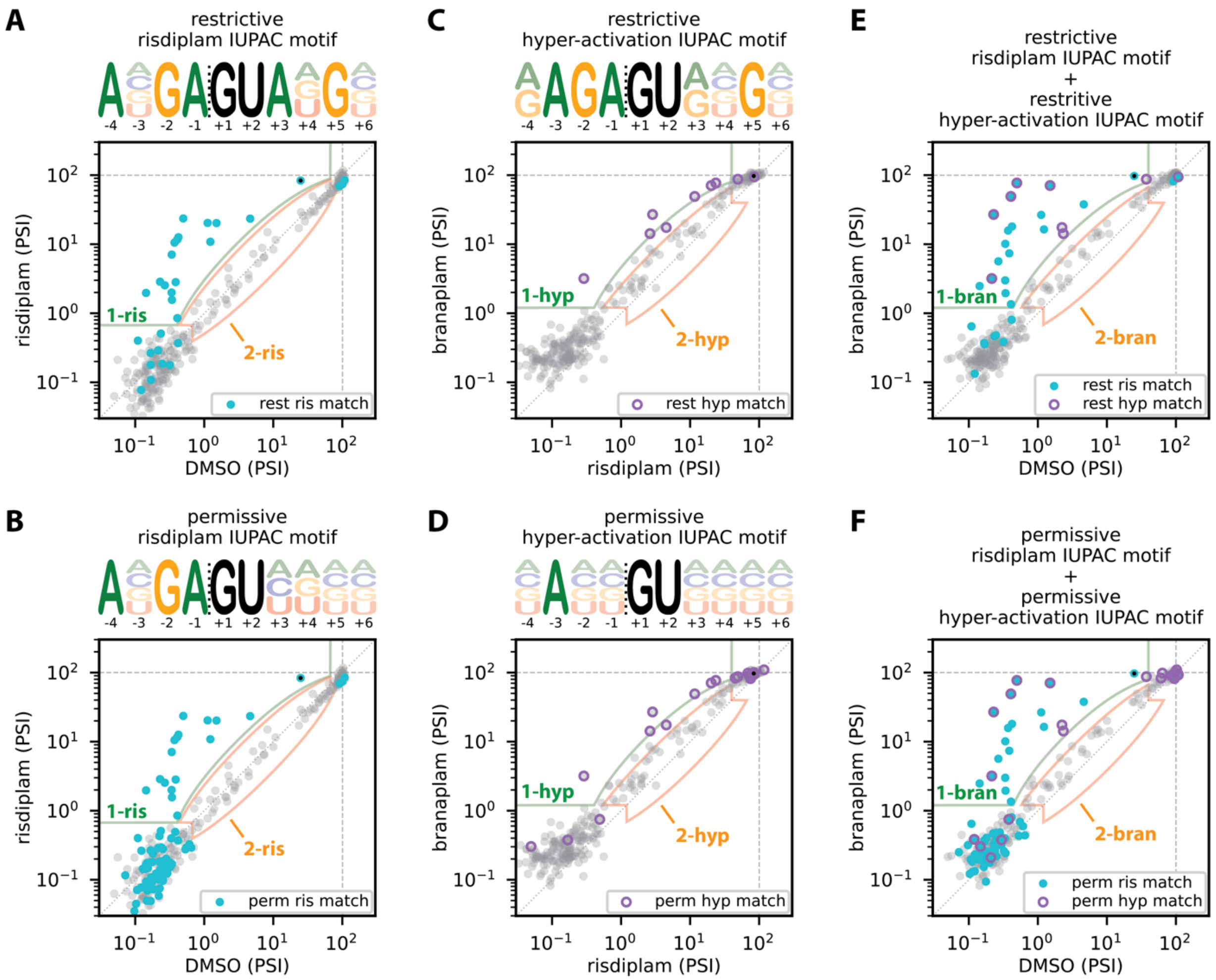

##### 6.3 *Figure S3. Multiple IUPAC motifs satisfy risdiplam vs. DMSO and branaplam vs. risdiplam classification criteria on* SMN2 *MPSA data*

(**A,B**) The (**A**) maximally restrictive and (**B**) maximally permissive IUPAC motifs that correctly classify 5'ss activated by (class 1-ris) or insensitive to (class 2-ris) risdiplam relative to DMSO. (**C,D**) The (**C**) maximally restrictive and (**D**) maximally permissive IUPAC motifs that correctly classify 5'ss that are activated by (class 1-hyp) or insensitive to (class 2-hyp) branaplam relative to risdiplam. (**E**,**F**) 5'ss that are activated by (class 1-bran) or insensitive to (class 2-bran) branaplam relative to DMSO are correctly classified by (**E**) the restrictive risdiplam IUPAC motif together with the restrictive hyper-activation IUPAC motif, as well as (**F**) the permissive risdiplam IUPAC motif together with the permissive hyper-activation IUPAC motif. MPSA, massively parallel splicing assay; 5'ss, 5' splice site; DMSO, dimethyl sulfoxide; PSI, percent spliced in.

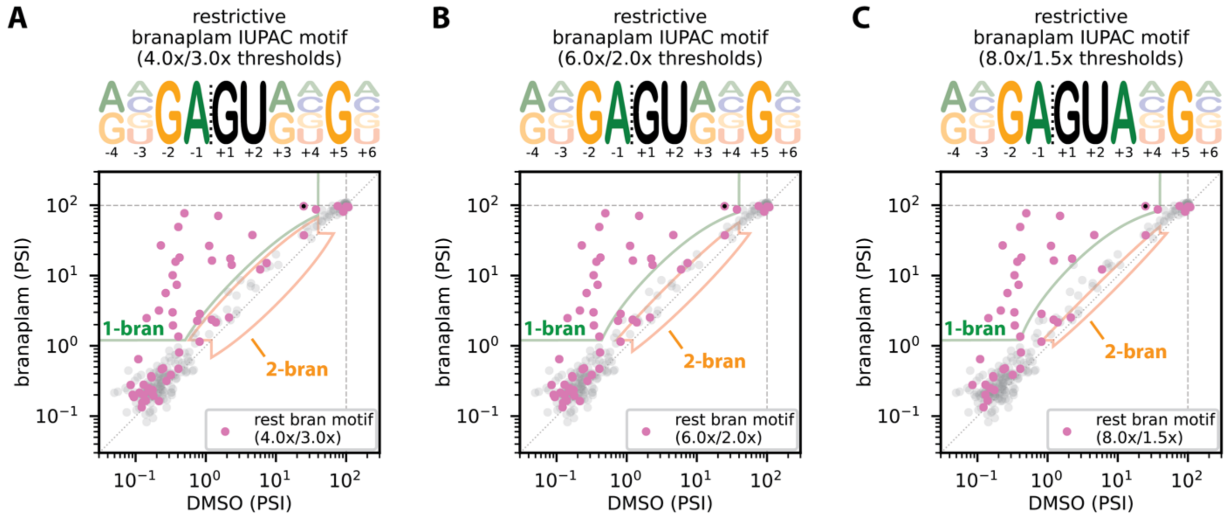

##### 6.4 *Figure S4. No single IUPAC motif satisfies branaplam vs. DMSO classification criteria on* **SMN2** *MPSA data*

All three panels (**A-C**) plot PSI values for 5’ss in the presence of branaplam versus DMSO as in **Fig. 1F**. Outlined areas show 5’ss classified as activated by branaplam (class 1-bran) or insensitive to branaplam (class 2-bran) using different fold-activation thresholds (indicated in panel titles): first number indicates the fold-activation limit on 5’ss in class 1-bran; second number indicates the fold-activation/repression limits on 5’ss in class 2-bran. Each panel also shows the restrictive IUPAC motif that matches all class 1-bran 5’ss, and highlights the assayed 5’ss that match this motif. Note that all three restrictive IUPAC motifs also match 5’ss that are insensitive to branaplam (i.e., are in class 2-bran), and thus no IUPAC motifs correctly classifies 5’ss based according to these definitions of class 1-bran and class 2-bran. Black dot, wild-type *SMN2* exon 7 5’ss (AGGA/GUAAGU). MPSA, massively parallel splicing assay. PSI, percent spliced in. 5’ss, 5’ splice site. DMSO, dimethyl sulfoxide.

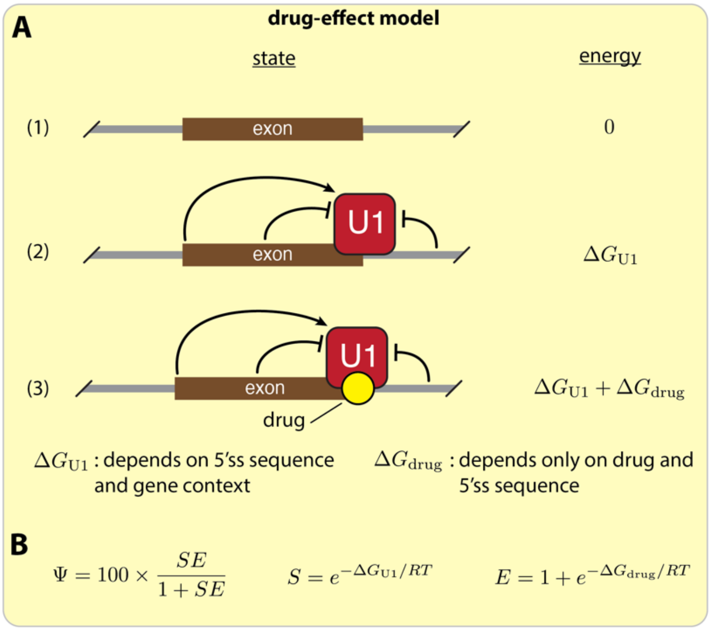

##### *6.5 Figure S5. Biophysical formulation of the allelic-manifold model*

(**A**) The allelic manifold model in **Fig. 2** follows from a thermodynamic model in which PSI is proportional to U1 occupancy on the 5'ss, the Gibbs free energy of U1 binding to the 5'ss sequence ($\Delta G_{\rm U1}$) depends on both 5'ss sequence and on gene context, and the Gibbs free energy of drug binding to the U1/5'ss complex ($\Delta G_{\rm drug}$) depends only on drug concentration and on 5'ss sequence. (**B**) The values of PSI ($\Psi$), context strength ($S$), and drug effect ($E$), written in terms of the Gibbs free energies from the thermodynamic model in **A**. See **Sec. 3.1** for details.

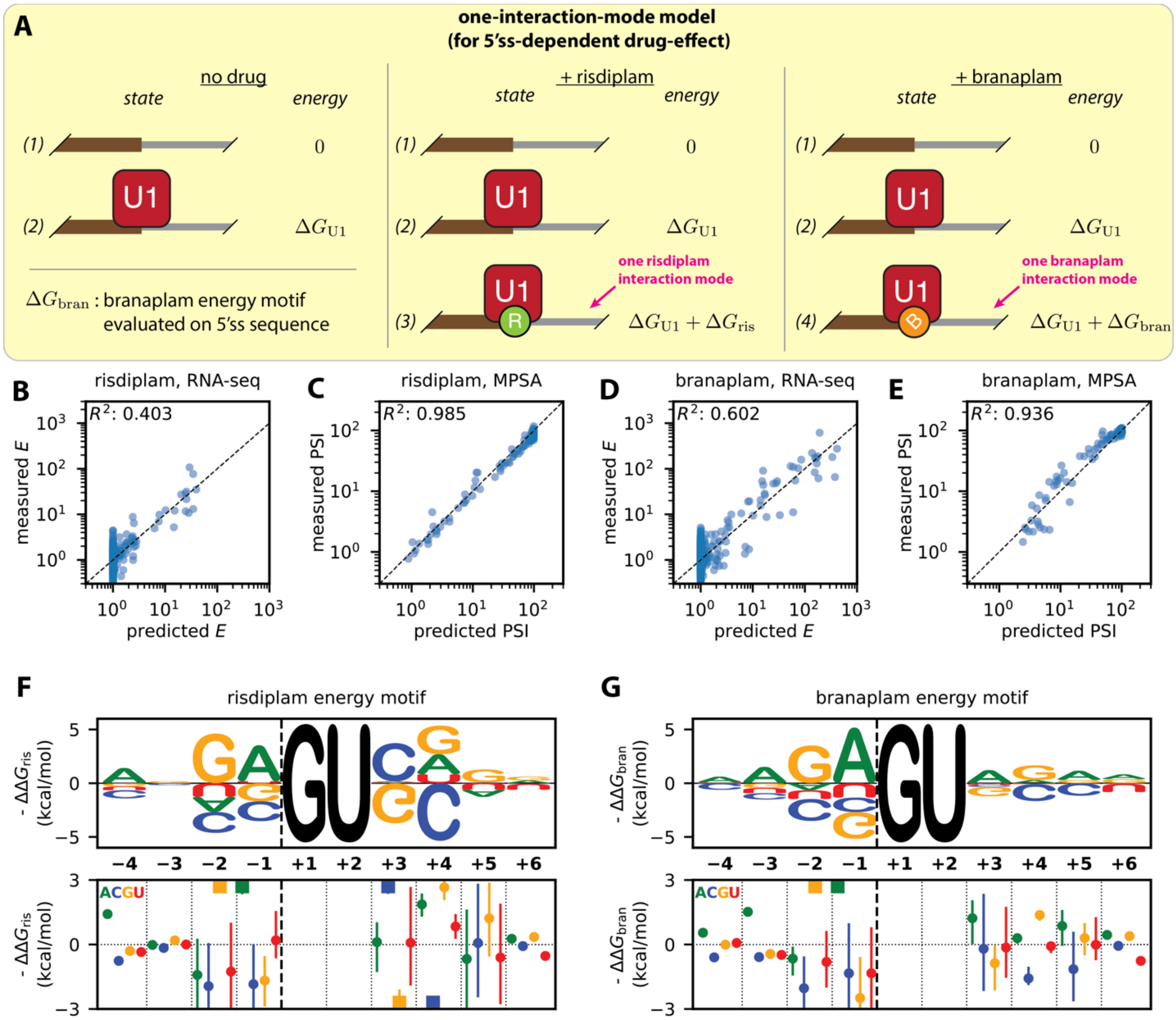

#### 6.6 *Figure S6. One-interaction-mode model for how risdiplam and branaplam affect splicing*

(**A**) PSI is assumed to be proportional to the equilibrium occupancy of U1 at the 5'ss. The model assumes three sequence-dependent Gibbs free energies: $\Delta G_{\mathrm{U1}}$, energy of U1 binding to the 5'ss; $\Delta G_{\mathrm{ris}}$, energy of risdiplam binding to the U1/5'ss complex; and $\Delta G_{\mathrm{bran}}$, energy of branaplam binding to the U1/5'ss complex. (**B-E**) Experimentally measured vs. model-predicted PSI values and drug-effect values based on the one-interaction-mode model. (**F,G**) Inferred additive parameters for (**G**) risdiplam binding energy and (**H**) branaplam binding energy. See **Fig. 3** legend for more annotation information. 5'ss, 5' splice site. MPSA, massively parallel splicing assay.

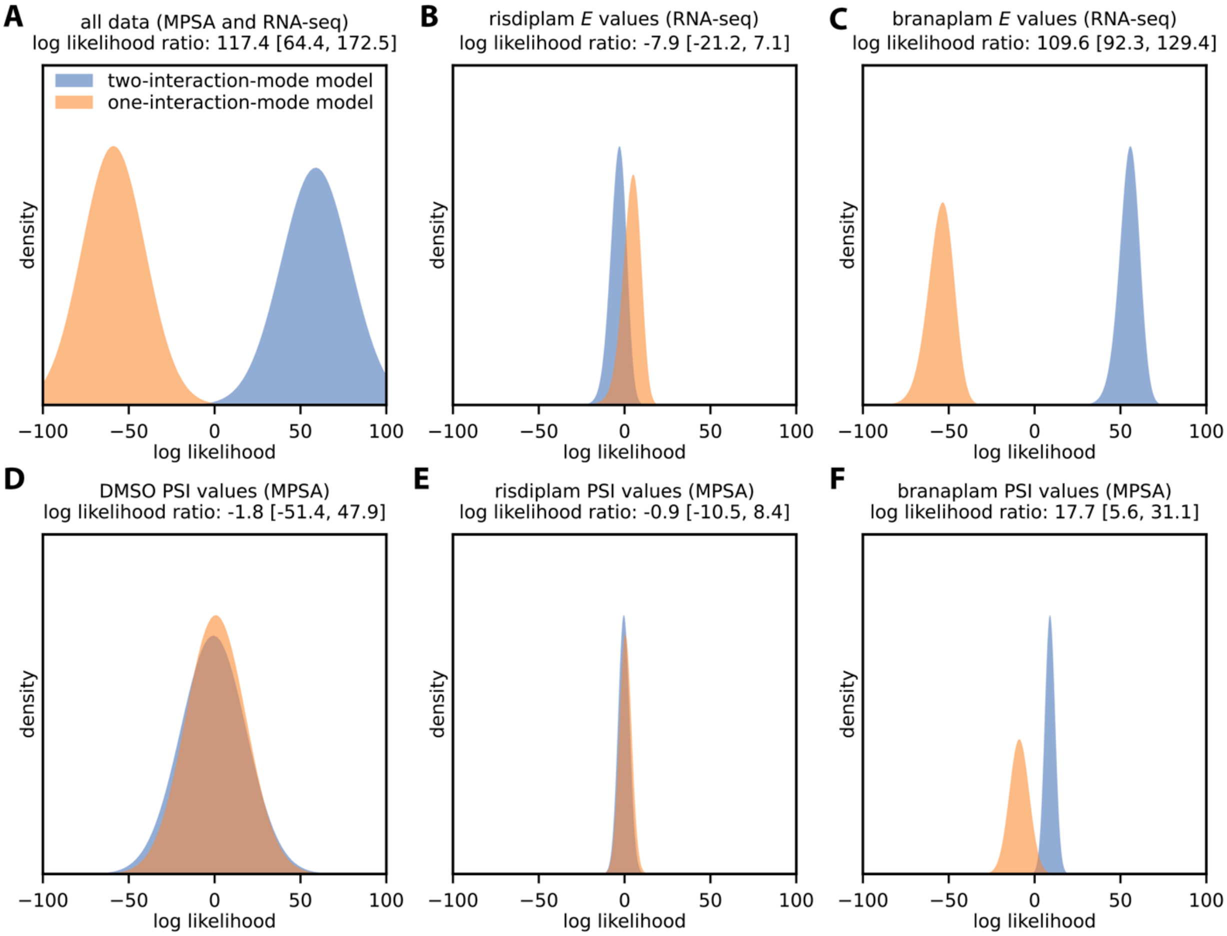

##### *6.7 Figure S7. Log likelihood values, total and stratified by dataset, for the two-interaction-mode model versus the one-interaction-mode model*

Shown are the distributions of log-likelihood values (centered about zero) for posterior-sampled parameters with log-likelihoods evaluated on: (**A**) all PSI values measured by MPSA and all drug effect values measured by RNA-seq, (**B**) drug effect values measured by RNA-seq for cells treated with risdiplam, (**C**) drug effect values measured by RNA-seq for cells treated with branaplam, (**D**) PSI values measured by MPSA for cells treated with DMSO, (**E**) PSI values measured by MPSA for cells treated with risdiplam, and (**F**) PSI values measured by MPSA for cells treated with branaplam. Probability distributions were estimated using DEFT[17]. PSI, percent spliced in. MPSA, massively parallel splicing assay.

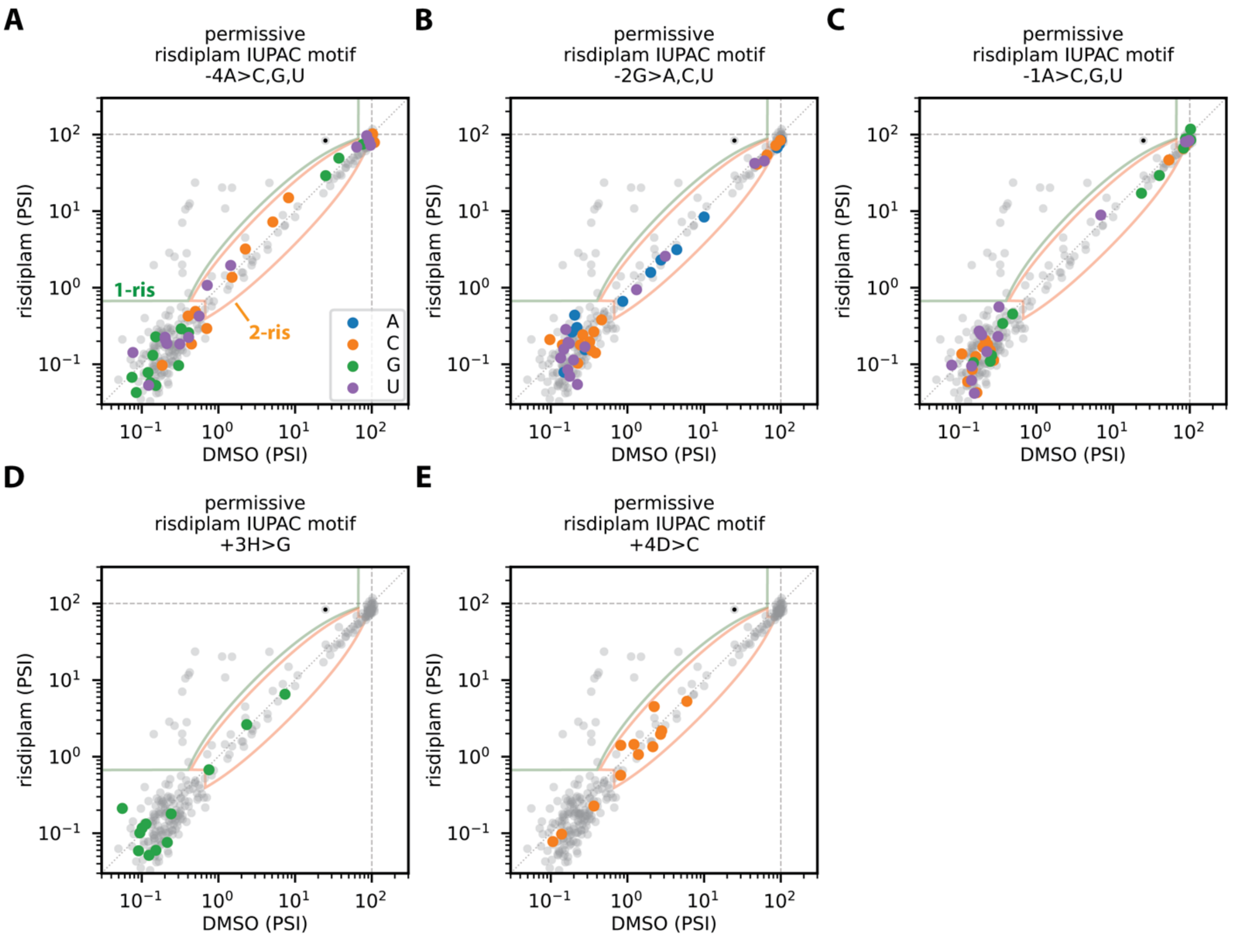

#### 6.8 *Figure S8. 5'ss substitutions that abrogate risdiplam activity*

(**A-E**) MPSA data for the *SMN2* exon 7 minigene library treated with risdiplam versus DMSO. Colored dots indicate 5'ss that match the risdiplam permissive IUPAC motif (ANGA/GUHDNN; **Fig. S3B**) except at a single position (compare to **Fig. S3B**). The mutations that lead to these single-nucleotide mismatches are: (**A**) A > C, G, or U at position -4; (**B**) G > A, C, or U at position -2; (**C**) A > C, G, or U at position -1; (**D**) A, C, or U > G at position +3; (**E**) A, G, or U > C at position +4. All of these mutations are seen to abrogate activation by risdiplam, since no 5'ss that have these mutations fall within the risdiplam-activation region (class 1-ris, green outlined area), and, for each of these mutations, some 5'ss fall within the risdiplam-insensitive region (class 2-ris, peach outlined area). 5'ss, 5' splice site. MPSA, massively parallel splicing assay. DMSO, dimethyl sulfoxide.

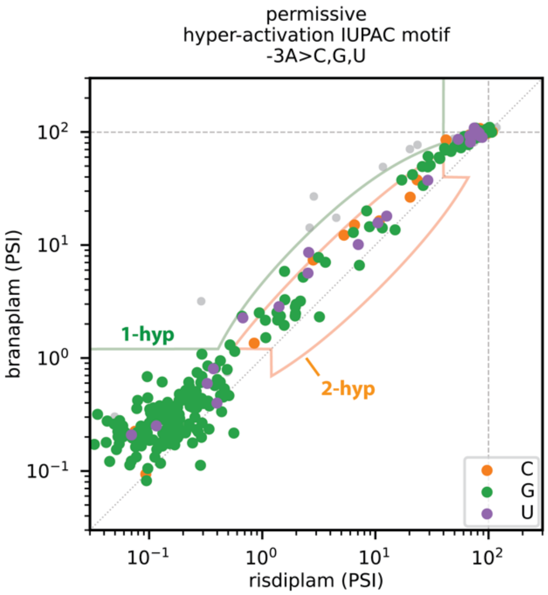

***6.9 Figure S9. 5’ss substitutions that abrogate hyper-activation by branaplam relative to risdiplam***
MPSA data for the *SMN2* exon 7 minigene library treated with branaplam versus risdiplam. Colored dots indicate 5’ss that do not have an A at position -3 and thus do not match the permissive hyper-activation IUPAC motif (NANN/GUNNNN; see **Fig. S3D**). All three mutations at this position (A > C, G, or U) are seen to abrogate hyper-activation, since no 5’ss that have these mutations fall within the hyper-activation region (class 1-hyp, light green outlined area), and, for each of these three mutations, some 5’ss fall within the no hyper-activation region (class 2-hyp, peach outlined area). 5’ss, 5’ splice site. MPSA, massively parallel splicing assay.

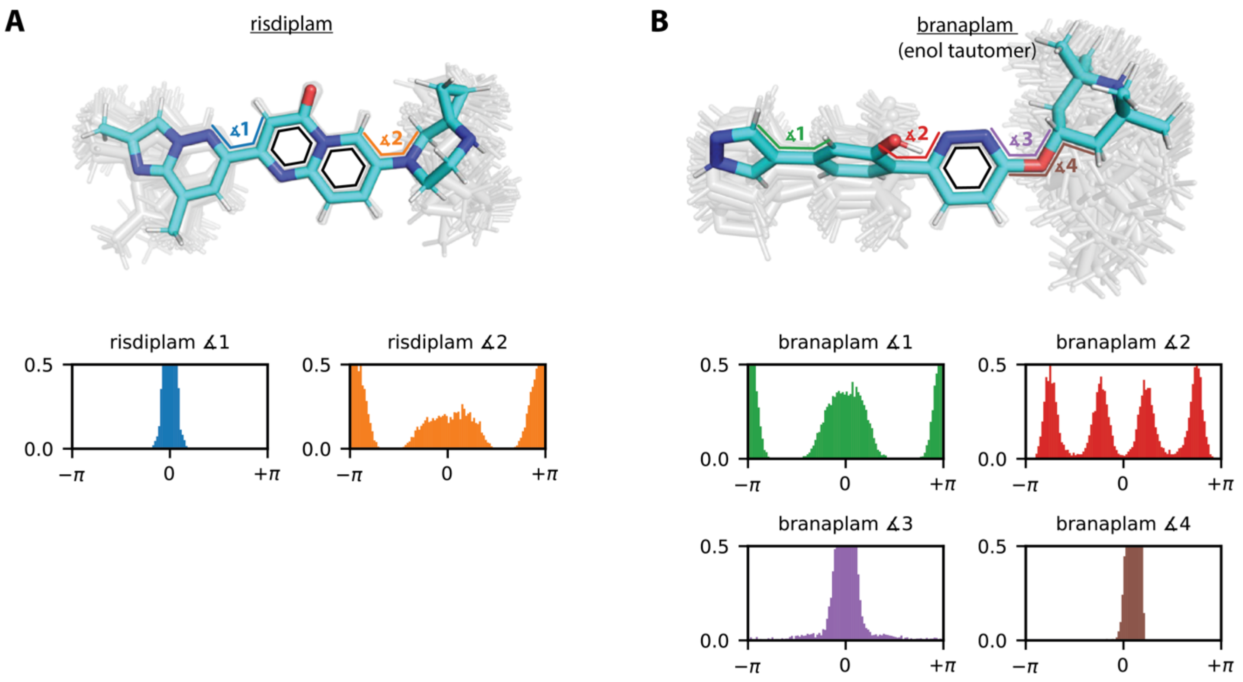

##### *6.10 Figure S10. Molecular dynamics simulations of free drug conformation*

Shown are molecular dynamics simulations for (**A**) risdiplam and (**B**) the enol tautomer of branaplam. Upper panels: starting conformation (cyan) with dihedral angles illustrated, together with structure clouds showing 100 conformations (grey) evenly sampled over a 1 µs simulation. Structures are aligned by the central carbon/nitrogen rings of each molecule (black hexagons). Lower panels: area-normalized histograms of the dihedral angles simulated for each molecule. Note that some distributions extend beyond the (shared) y-axis limits.

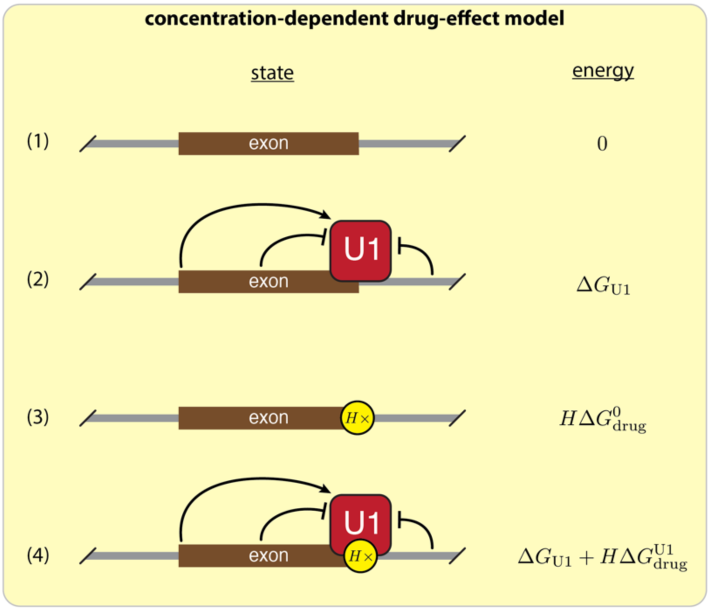

##### *6.11 Figure S11. Biophysical formulation of the concentration-dependent drug-effect model*

This model assumes a 5’ss can be in one of four possible states: (1) not bound by U1 or drug, (2) bound by U1, (2) bound by $H$ molecules of drug, or (4) bound by U1 and $H$ molecules of drug. $\Delta G_{\mathrm{U1}}$, Gibbs free energy of U1 binding to a 5’ss. $\Delta G^{0}_{\mathrm{drug}}$, Gibbs free energy of a single drug molecule binding to the 5’ss in the absence of U1. $\Delta G^{\mathrm{U1}}_{\mathrm{drug}}$, Gibbs free energy of a single drug molecule binding to the U1/5’ss complex. $H$, Hill coefficient. See Supplemental Information Text for formulas that relate these quantities to those in **Fig. 5C,D** (i.e., $[\mathrm{drug}]$, $S$, $\mathrm{EC}_{2\mathrm{x}}$, and $E_{\mathrm{max}}$).

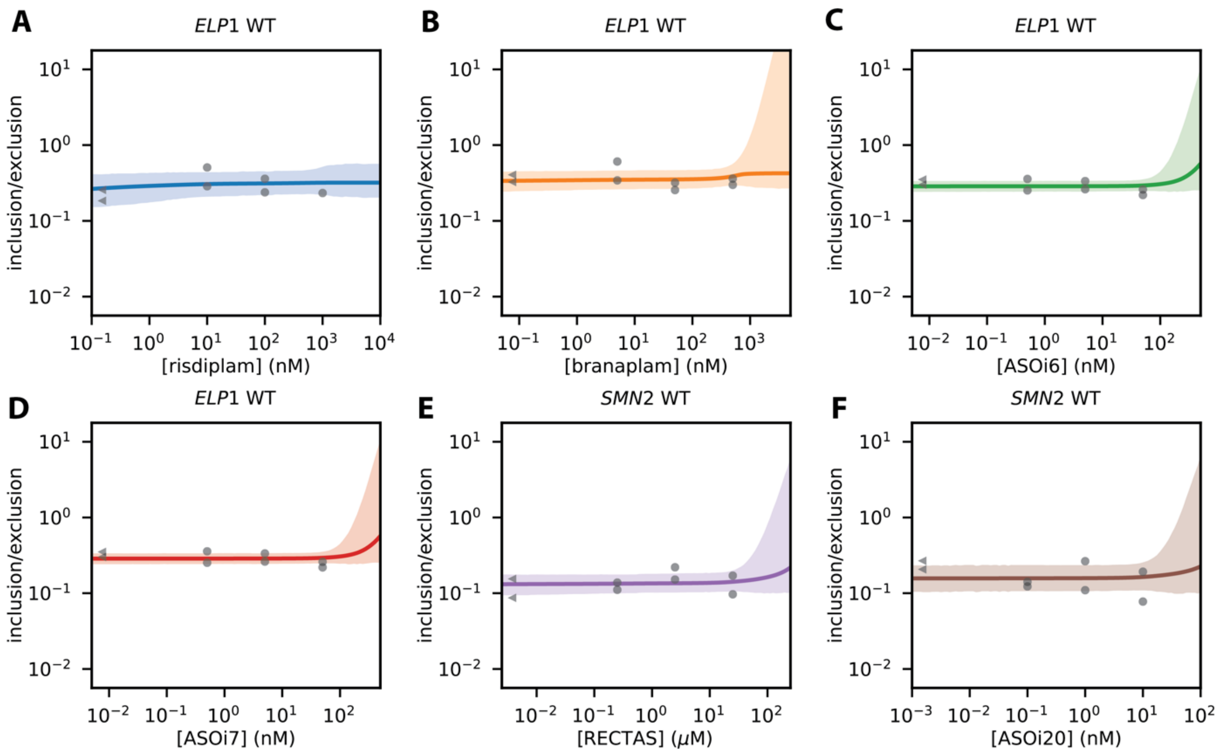

#### *6.12 Figure S12. Dose-response curves for negative control minigenes*

Single-drug dose-response data and inferred curves for splice-modifying drugs on non-target minigenes. (**A-D**) *ELP1* exon 20 minigene in response to (**A**) risdiplam, (**B**) branaplam, (**C**) ASOi6, and (**D**) ASOi7. (**E,F**) *SMN2* exon 7 minigene in response to (**E**) RECTAS and (**F**) ASOi20. Each panel shows data from 4 drug concentrations. Dots show biological replicates (median of 3 technical replicates). Curves and shaded regions show the results of Bayesian inference using the same model as in **Fig. 5E-L** and **Fig. 6A-D**.

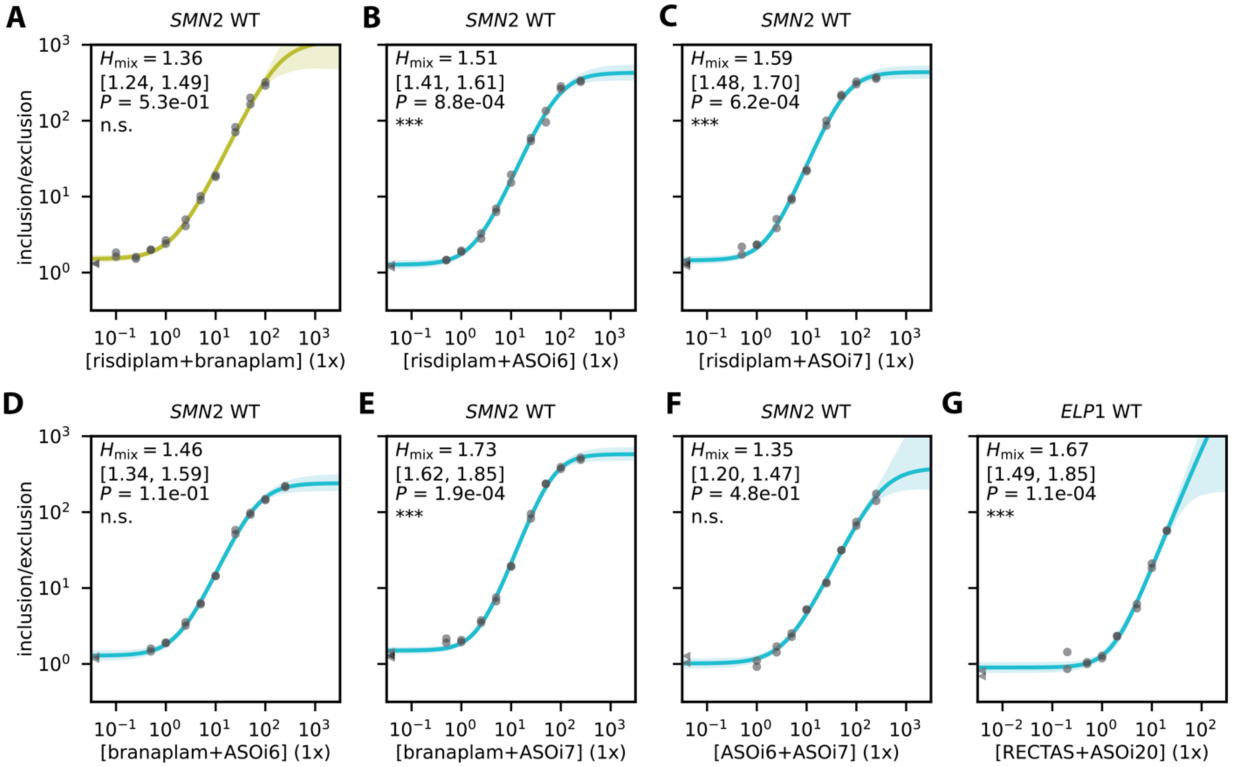

##### *6.13 Figure S13. Multi-drug synergy among splice-modifying drugs determined from dose-response data*

In addition to the two-drug linear mixture data presented in Fig. 6E-K, we assessed the presence or absence of synergy between pairs of drugs based on whether the Hill coefficient of a two-drug cocktail was larger than the Hill coefficients of the two individual drugs in the cocktail. (**A-G**) Two-drug dose-response curves for *SMN2* exon 7 in response to (**A**) a risdiplam/branaplam cocktail, (**B**) a risdiplam/ASOi6 cocktail, (**C**) a risdiplam/ASOi7 cocktail, (**D**) a branaplam/ASOi6 cocktail, (**E**) a branaplam/ASOi7 cocktail, and (**F**) an ASOi6/ASOi7 cocktail. (**G**) Two-drug dose-response curve for *ELP1* exon 20 in response to a RECTAS/ASOi20 cocktail. A 1x cocktail corresponds to a mixture of 0.5x of each component drug, where the 1x concentration (corresponding to $\mathrm{EC}_{2x}$ values measured in preliminary experiments) is 14 nM for risdiplam, 7 nM for branaplam, 0.1 nM for ASOi7, 0.6 nM for ASOi6, 300 nM for RECTAS, and 0.08 nM for ASOi20. $H$, single-drug Hill coefficient (median and 95% credible interval). $H_{\mathrm{mix}}$, drug cocktail Hill coefficient (median and 95% credible interval). $P$, p-value for no-synergy null hypothesis (i.e., that $H_{\mathrm{mix}}$ is not larger than both $H_{\mathrm{drug\,1}}$ and $H_{\mathrm{drug\,2}}$) computed using Hamiltonian Monte Carlo sampling. **Table S1** summarizes the Hill coefficients and p-values for the relevant dose-response curves.

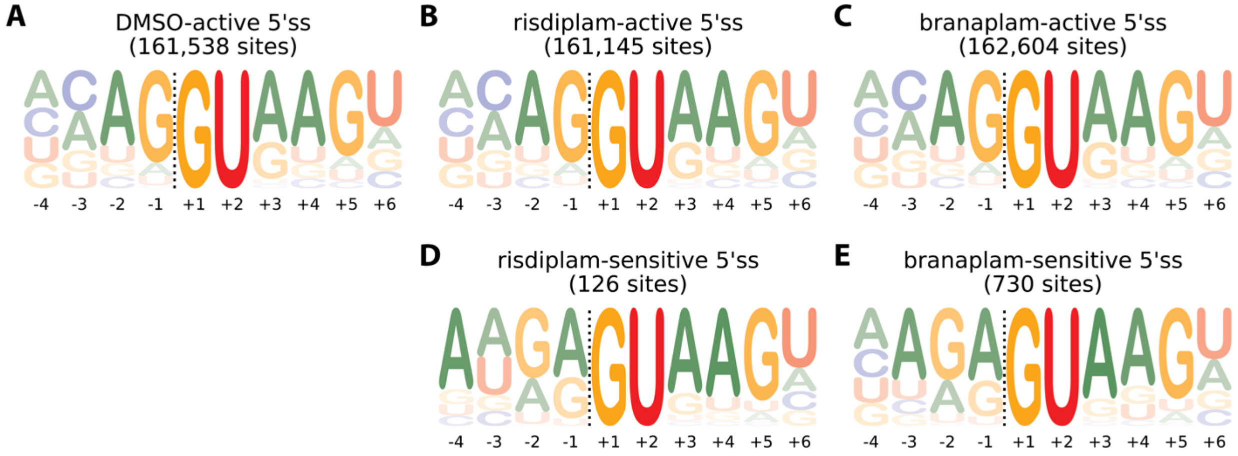

***6.14 Figure S14. Probability logos derived from RNA-seq data.***

Probability logos[18] representing sets of 5’ss sequences present in the human genome and satisfying various criteria on PSI as measured by RNA-seq. (**A**) 5’ss active in DMSO-treated cells ($PSI_{DMSO}$ > 10). (**B**) 5’ss active in risdiplam-treated cells ($PSI_{ris}$ > 10). (**C**) 5’ss active in branaplam-treated cells ($PSI_{bran}$ > 10). (**D**) 5’ss sensitive to risdiplam ($PSI_{ris}$ > 10, $PSI_{DMSO}$ < 90, $\text{logit}_2\ PSI_{ris}$ - $\text{logit}_2\ PSI_{DMSO}$ > 4). (**E**) 5’ss sensitive to branaplam ($PSI_{bran}$ > 10, $PSI_{DMSO}$ < 90, $\text{logit}_2\ PSI_{bran}$ - $\text{logit}_2\ PSI_{DMSO}$ > 4). $\text{logit}_2\ \text{PSI} = \log_2 \left[\left(\frac{\text{PSI}}{100}\right) / \left(1 - \frac{\text{PSI}}{100}\right)\right] = \log_2 SE$ where $S$ denotes context strength and $E$ denotes drug effect, and thus $\text{logit}_2\ \text{PSI}_{\text{drug}} - \text{logit}_2\ \text{PSI} = \log_2 E_{\text{drug}}$. Note that the logos illustrating 5’ss active in the presence of risdiplam and branaplam (panels **B** and **C**) are nearly identical to the logos illustrating 5’ss active in the presence of DMSO (panel **A**) and do not show hallmarks of drug-dependent activity. The logos illustrating 5’ss activated by risdiplam or branaplam (panels **D** and **E**) show do some sequence patterns present in the interaction-mode-specific IUPAC motifs (**Fig. 1**) and energy motifs in (**Fig. 3**)—e.g., the enrichment of $A_{-4}N_{-3}G_{-2}A_{-1}$ among 5’ss activated by risdiplam, and the enrichment of $N_{-4}A_{-3}G_{-2}A_{-1}$ among 5’ss activated by branaplam. However, the logos in panels E and E convolve these sequence patterns with drug-independent sequence patterns present in the DMSO probability logo (panel **A**).
